# Precision design of single and multi-heme de novo proteins

**DOI:** 10.1101/2020.09.24.311514

**Authors:** George H. Hutchins, Claire E. M. Noble, Hector Blackburn, Ben Hardy, Charles Landau, Alice E. Parnell, Sathish Yadav, Christopher Williams, Paul R. Race, A. Sofia F. Oliveira, Matthew P. Crump, Christiane Berger-Schaffitzel, Adrian J. Mulholland, J. L. Ross Anderson

## Abstract

The *de novo* design of simplified porphyrin-binding helical bundles is a versatile approach for the construction of valuable biomolecular tools to both understand and enhance protein functions such as electron transfer, oxygen binding and catalysis. However, the methods utilised to design such proteins by packing hydrophobic side chains into a buried binding pocket for ligands such as heme have typically created highly flexible, molten globule-like structures, which are not amenable to structural determination, hindering precise engineering of subsequent designs. Here we report the crystal structure of a *de novo* two-heme binding “maquette” protein, 4D2, derived from the previously designed D2 peptide, offering new opportunities for computational design and re-engineering. The 4D2 structure was used as a basis to create a range of heme binding proteins which retain the architecture and stability of the initial crystal structure. A well-structured single-heme binding variant was constructed by computational sequence redesign of the hydrophobic protein core, assessed by NMR, and utilised for experimental validation of computational redox prediction and design. The structure was also extended into a four-heme binding helical bundle resembling a molecular wire. Despite a molecular weight of only 24kDa, imaging by CryoEM illustrated a remarkable level of detail in this structure, indicating the positioning of both the secondary structure and the heme cofactors. The design and determination of atomic-level resolution in such *de novo* proteins is an invaluable resource for the continued development of novel and functional protein tools.

## Introduction

Computational, *de novo* protein design has attained a level of sophistication where atomistic precision is almost routine^1,2^, and there now exist a multitude of examples demonstrating our command over these fundamental biomolecular building blocks^3,4^. Conversely, where oxidoreductase cofactors have been successfully incorporated into simplified protein scaffolds^5,6,7^, principally termed maquettes, there are limited examples where high-resolution structural information has been successfully obtained^8,9,10^. This shortfall hinders the downstream engineering of these robust and versatile scaffolds to incorporate substrate binding sites, tailor active site residues and fine tune cofactor biophysical properties in a predictable manner. Ultimately, such fine control of structure will lead to significant improvements in these proteins, aiding the expansion of their functional and catalytic repertoire, and enabling, for instance, imprinted regio- and stereoselectivity in *de novo* oxidoreductase enzymes.

Despite the drive toward well-packed, native-like states in *de novo* proteins, it should be noted that certain heme-containing maquettes exhibit catalytic activities comparable to their natural counterparts while adopting conformationally dynamic structures more reminiscent of molten globules than well-folded native-like states^11,12^. In these cases, the dynamic nature of the protein may in fact enhance catalytic activity by lowering barriers to substrate entry and product exit, though the relationship between dynamics and catalysis in these simple proteins currently remains unclear^11,13,14^. Since these activities extend to industrially and biosynthetically valuable reactions^12^, it would be prudent to explore this relationship in greater depth and establish a framework of robust, engineerable *de novo* proteins to address these and other fundamentally important questions relating to biologically relevant phenomena, such as electron transfer.

To this end, here we describe the design and construction of single and multi-heme proteins with well-defined structures and biophysical properties that can be predictably fine-tuned. Our strategy was based on the successful design and characterisation of the D2 peptide by the Degrado lab^15^, which self assembles into a diheme tetrahelical bundle. Following the addition of simple loops, we created a single chain variant with nanomolar heme affinity, 4D2, that we were able to crystalise, obtaining a high-resolution structure of the heme-bound maquette. This structure guided the subsequent computational design of a rigid monoheme maquette, and an extended tetraheme maquette. We obtained further structural insight into our designs using NMR spectroscopy and cryo-electron microscopy, the latter enabling heme-enhanced visualization of our 24 kDa tetraheme maquette which represents the current mass limit of the technique. This structural information and our confidence in the fidelity of the design process enabled the implementation of Monte Carlo Continuum Electrostatic Calculations^16^ to produce further maquettes with predictably altered redox potentials. Such fine-tuning of a fundamental biophysical property of the cofactor is central to heme protein engineering and the future construction of efficient oxidoreductases on our own terms.

## Results

### Conversion of the D2 peptide to an in vivo expressed single-chain protein

For a design process with higher precision than that used in the construction of the earlier maquettes, we selected the D2 peptide^15^ reported by Ghirlanda *et al,* as a starting point for further design. D2 was originally designed by parameterizing the transmembrane cytochrome *b* subunit of the cytochrome bc1 complex, using computational methods to define a sequence that would self-assemble into a soluble tetrahelical bundle with overall D2 symmetry in the presence of heme. The resulting peptide demonstrated high (but undefined) affinity for two heme Bs within the assembly, and relatively well-resolved 2D HSQC NMR spectra were observed in presence of the heme B cofactors. Despite these promising observations, no high-resolution structural information was obtained. We reasoned that the heme B binding affinity could be improved by creating a single chain tetrahelical bundle, potentially preorganising the heme binding sites and reducing the entropic cost of complex assembly in the unconnected D2:heme B assembly. Given the computational design of the heme binding sites and NMR data, it was also hoped that the single chain variant would retain the apparent singular, near-native structure of the original assembly.

To achieve this single chain variant of D2, we designed a protein where four copies of the 25-residue D2 peptide were linked together by three short loops: two TSN loops between helices 1-2 and 3-4; and one GSVSP sequence at the central loop between helices 2-3. We also included a TEV (Tobacco Etch Virus N1A) protease-cleavable hexahistidine tag and V5 epitope at the N-terminus to enable purification by metal affinity chromatography and antibody detection respectively, resulting in a 112-amino acid four helix bundle after TEV-cleavage. Following creation of a synthetic gene and expression in T7 Express (NEB) *E. coli* from pET151 (Thermo), a vibrantly red cell pellet was obtained, indicating *in vivo* heme B binding (Fig. 1). Whilst cytoplasmic expression of the protein with or without supplementation with the heme precursor γ-aminolevulinic acid^17^ can yield significant heme binding, the reliability of cofactor incorporation was increased considerably by periplasmic export, facilitated by cloning 4D2 into a modified pMal-p4x vector (pSHT) containing a cleavable, N-terminal signal sequence for periplasmic export, used previously for the expression of *de novo* c-type maquettes^18^. Low heme loaded cytoplasmic preparations can be recovered either through titration of hemin into purified protein, or through supplementation with excess hemin on cell lysis. Reconstitution of 4D2 with heme B results in protein with identical biophysical and spectroscopic properties to periplasmically-expressed 4D2 or cytoplasmically-expressed 4D2 with a full heme complement. In addition, to facilitate heme B binding studies and to assess the biophysical characteristics of the apo-4D2, we were able to remove bound heme B using acid:2-butanone extraction, as previously described^19^.

**Figure 1:**
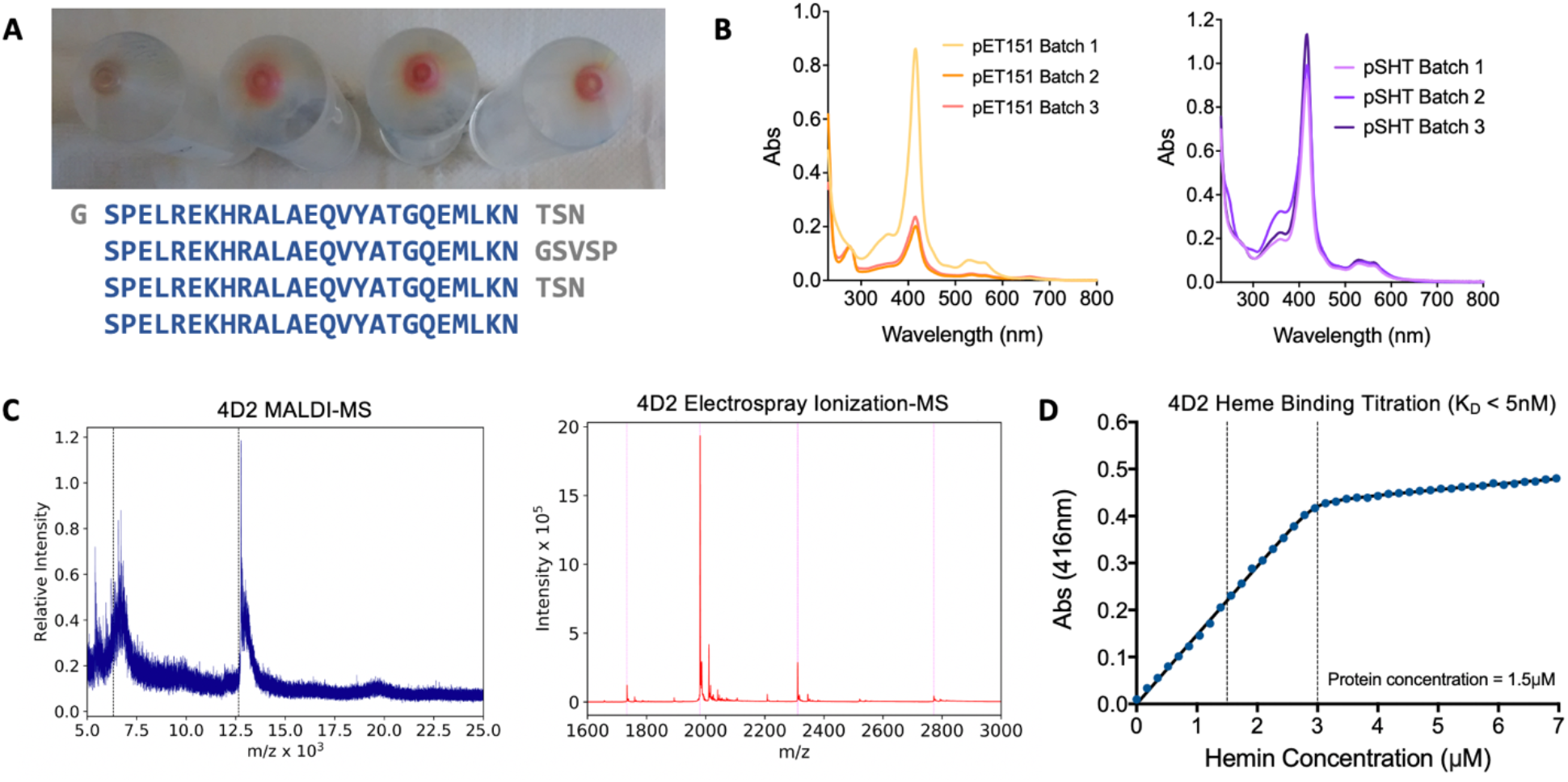
The single chain polypeptide 4D2 can be fully assembled with heme by expression in E. coli, although heme binding is variable – all four growth conditions in the representative cell pellets pictured (A) are identical but were cultured from different colonies from the same transformation. The protein was expressed in all four cultures, but minimal heme incorporation is observed from one of the colonies (left). Reliability of heme binding in vivo was improved by expression with a periplasmic export tag in the pSHT vector (B) compared to cytoplasmic expression in pET151. Mass spectrometry (C) confirms the molecular weight of the apo protein by MALDI (12.63kDa) and the assembled two-heme bound holo protein by ESI-MS (13.87 kDa, m/z peaks correspond to Z=5, 2773.02, Z=6 2208.10, Z=7 1980.85 and Z=8 1733.14). Binding titrations further demonstrate tight binding of heme in a 2:1 ratio.

Heme-bound 4D2 exhibits UV/visible spectra typical of proteins binding heme B by bis-histidine ligation, with a distinctive Soret peak at 416 nm in the oxidized, ferric state^5,20,21^. We subsequently used electrospray ionization mass spectrometry (ESI-MS) under aqueous, non-denaturing conditions^22^ to confirm the protein mass and further examine heme binding. Under these soft ionization conditions, we principally observe the diheme 4D2 complex at the correct mass, with only with only a small proportion of monoheme- or apo-4D2 m/z peaks observed (Figure 1C). Conversely, the heme groups dissociate under the conditions of MALDI mass spectrometry, resulting solely in the detection of apo-4D2. Following titrations of hemin into apo-4D2, we established tight heme B binding, with an observed dissociation constant (K_D_) of < 5 nM, and with no evidence of negative cooperativity. Using circular dichroism, we also observe that heme binding dramatically increases the helicity and thermal stability of 4D2; apo-4D2 is fully unfolded at 37°C, while diheme 4D2 demonstrates only a small loss in helicity until the start of a more cooperative melting event above 80°C (Fig. 2). To determine the heme redox potentials, we used optically transparent thin layer (OTTLE) electrochemistry^23^, observing midpoint potentials of −182 mV and −115 mV, and, in notable contrast to the equivalent data obtained for diheme D2, there is no evidence of hysteresis in these redox titrations. Also, unlike the original D2 peptide complex^15^, the 2D H^1^-N^15^-HSQC spectra of diheme 4D2 exhibited only moderate signal dispersion (Supplementary Fig. 1), offering only minimal potential of peak or structural assignment by NMR. This could indicate that multiple protein conformations exist and possibly interconvert on the NMR timescale, or possibly that there exists another source of structural heterogeneity in diheme 4D2.

**Figure 2:**
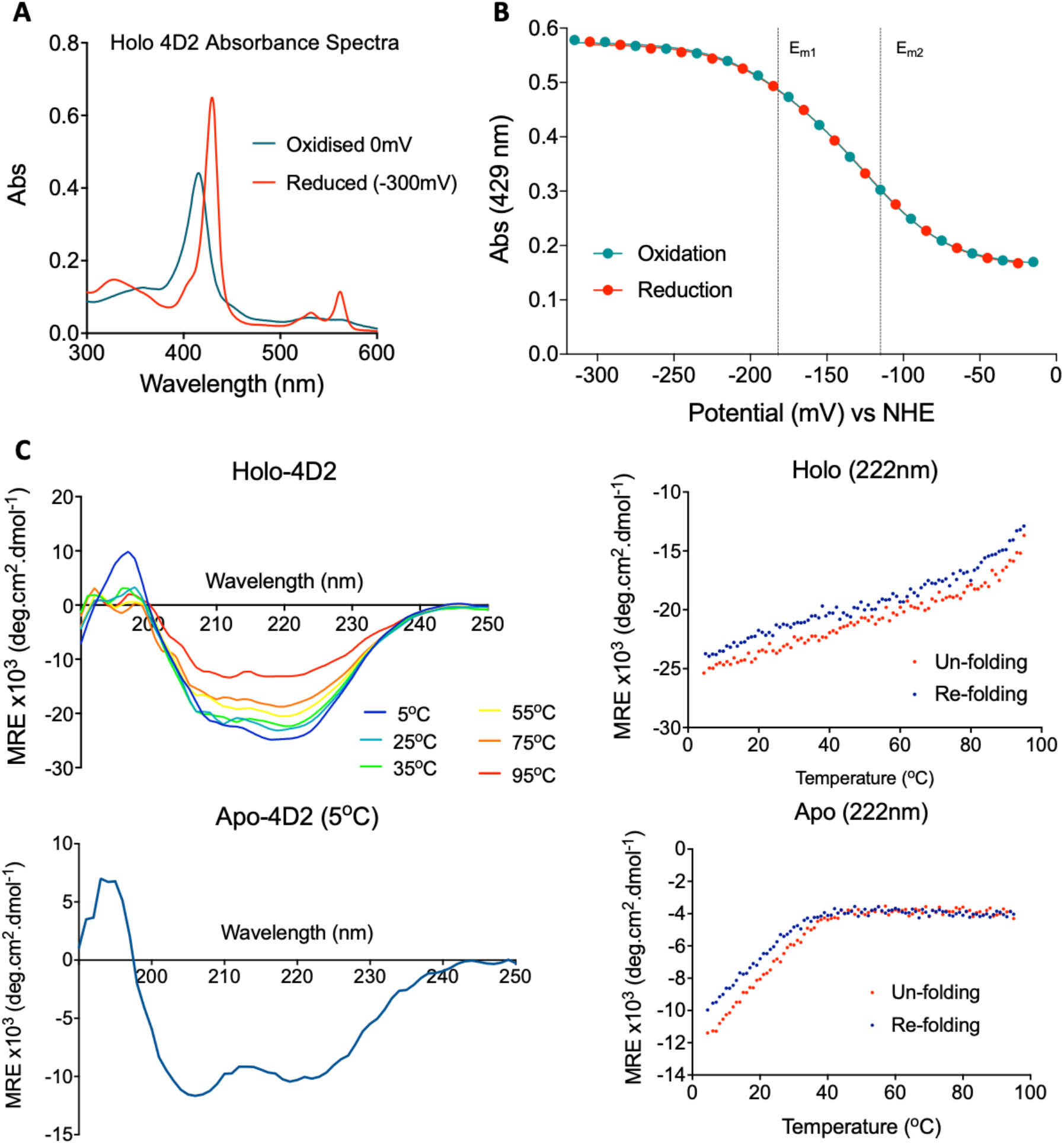
(A) The characteristic absorbance spectra of ligated b-type heme shifts upon reduction from 416nm to 429nm at the Soret peak alongside the expected splitting of the Q-bands. (B) Redox potentiometry measured using an OTTLE (optically transparent thin layer electrochemistry) cell demonstrated reversible reduction of the protein, fitted to a 2×1e-Nernst equation to determine the midpoint potential of the two sequential electron transfers. (C) Far-UV Circular dichroism spectroscopy indicated the helicity and thermostability of the holo protein, in comparison to a relatively unstructured and unstable apo-form.

### Crystal Structure of a de novo Heme-Bound Maquette

Though the NMR indicated structural heterogeneity, we successfully obtained crystals of the diheme 4D2 from 96-well sitting drop screening plates, though crystal formation was slow, requiring 4 months for the appearance of the first crystals and 6 months to produce diffraction-quality crystals. We were able to collect datasets at four wavelengths at the I03 beamline at Diamond light source, the fluorescence profile of the two iron atoms of the heme groups facilitating experimental phase determination by multi-wavelength anomalous dispersion (MAD)^24^. We subsequently determined the crystal structure to a resolution of 1.9 Å (Fig. 3), representing one of the first reported structures of a heme- or porphyrin-binding *de novo* protein.

**Figure 3:**
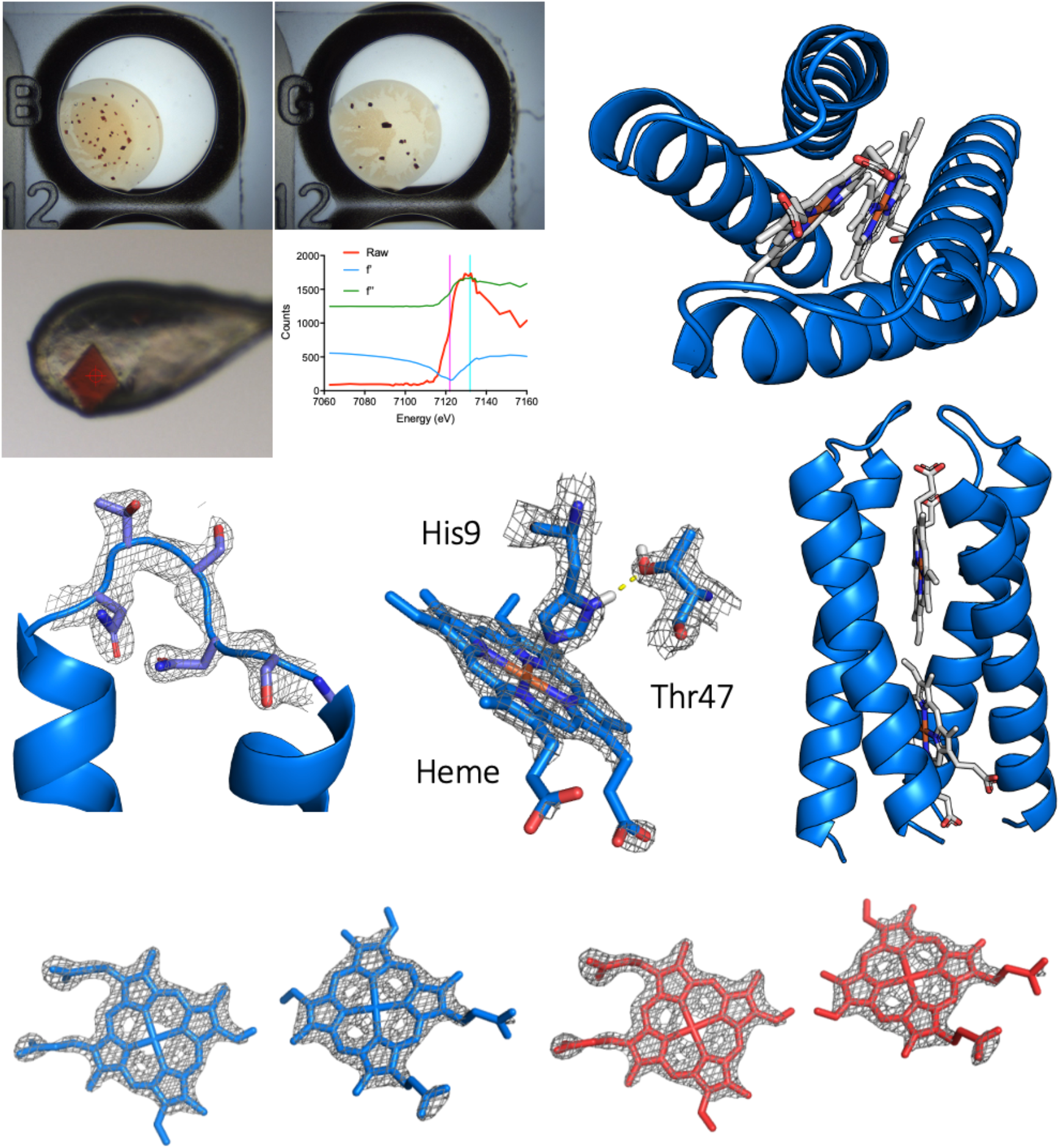
4D2 formed diffraction quality crystals but growth required up to six months of vapour drop diffusion, whilst the two heme-irons enabled experimental phasing by MAD using wavelengths determined by a fluorescence edge scan. The structure conforms to D2 symmetry around the ligated heme groups, connected by three loops, two of which are sufficiently rigid to be resolved by crystallography. One of the four potential hydrogen bonding interactions between the ligating histidine residues and a nearby “keystone” threonine has also been highlighted. Ambiguous electron density suggested that the heme groups likely bind in multiple orientations, lacking specificity to position the asymmetric vinyl groups. Two of the four potential orientations are illustrated, with the vinyl groups either opposing (blue) or adjacent (red) based on a flipping of the porphyrin ring.

**Figure 4:**
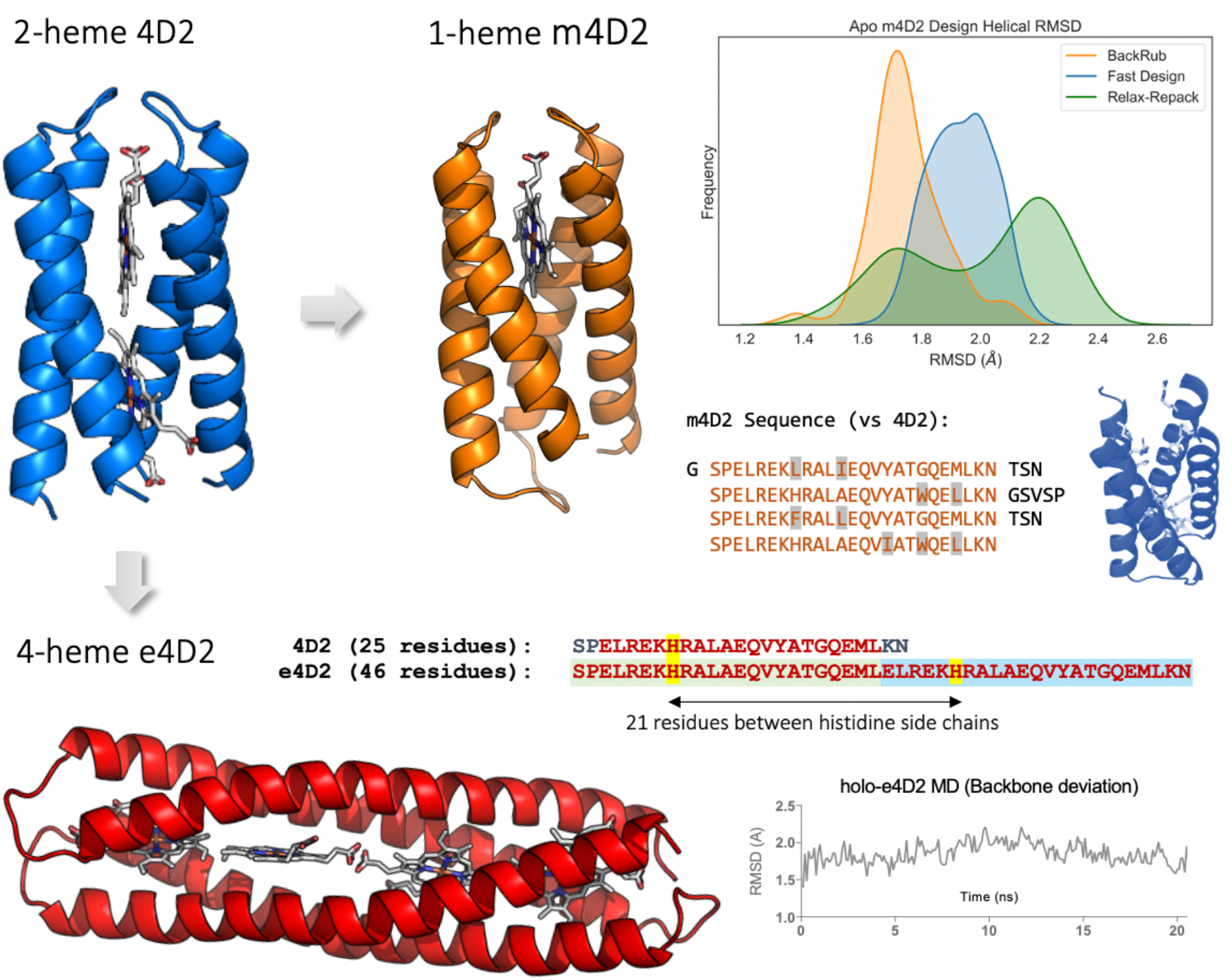
The crystal structure of 4D2 was used as a starting model to design single-heme and four-heme variants, increasing the diversity of subsequent potential design strategies. The sequence and structure of the hydrophobic core in the single heme binding m4D2 was redesigned an optimised using a series of simple Rosetta design scripts (included in Supplementary Information), and molecular dynamics simulations were used to identify stable, single conformation structures in the apo form using RMSD and RMSF (data not shown) analysis of 100 ns trajectories. The initial model of e4D2 was designed by building extended coiled-coil structures using ISAMBARD fitted to the 4D2 crystal structure, docking the heme groups, and minimizing the structure using a combination of Rosetta and MD.

The solved crystal structure of 4D2 matches very well to the expected fold, with four helices arranged in an ordered coiled coil-like structure and the four histidine residues ligating each of the two heme cofactors across opposite helices. The identical helical regions fit the D2 symmetry of the original parameterized peptide design^15^. Each histidine is contacted by a threonine within an approximate hydrogen bonding distance, most likely forming the ‘keystone’ hydrogen bonding interactions in the original design. While the two shorter TSN loops on one side of helical bundle are resolved, the flexible GSVSP loop between helices two and three is not observed in this structure, suggesting a relatively disordered region around the adjacent protein termini.

Interestingly, there is evidence of disorder in the heme B orientations within the binding pockets, with extended density visible in positions 1, 2, 3 and 4 of the tetrapyrrole ring. We attribute this to the presence of two binding modes, related to each other by a 180° rotation of the asymmetric heme, placing the 2 and 4 vinyl groups in the apparent 1 and 3 methyl positions of the corresponding other orientation. This has been observed in other heme B binding proteins, including neuroglobin^25^ and several bacterioferritins^26,27^, where near equal occupancy of the two modes was observed. This may also explain the poor signal dispersion in the NMR; combined with the repetitive sequence of the four helices, the four possible combinations of heme orientations would certainly lead to significant peak broadening and relatively poor dispersion. So, while heterogeneity seems apparent from the NMR data, it may not be due to the global conformational dynamics of the protein, and the diheme 4D2 may possess more native like structure in solution than was initially presumed. The edge-to-edge distance between the heme cofactors is small (2.6 Å) and close to being within Van der Waals contact, facilitating rapid electron transfer between the heme groups^28^.

### Expansion and Redesign: Single and Multi-Heme Variants of 4D2

Following the successful determination of the 4D2 structure, we reasoned that the scaffold could provide a template for further heme protein design, enabling us to access single and multi-heme proteins with similar atomistic control. We initially designed a monoheme variant, m4D2, using Rosetta^29^ to remove one of the heme binding sites and repack the vacated binding pocket in the protein core. We selected the heme binding site adjacent to the termini and longer GSVSP loop for removal, and used a flexible backbone design protocol^30^ to mutate key positions in the core to an abbreviated library of hydrophobic amino acids. To minimize unnecessary and potentially destabilizing changes to the protein, we used SOCKET^31^ to identify key knobs-into-holes interactions, and avoided modification of the residues involved in these contacts. In total, we selected 11 residues for the redesign process, representing about 10% of the total protein, of which, 9 were mutated in the final sequence of m4D2. The flexible backbone protocol we employed utilized the Backrub method^32^ for backbone sampling, applying relatively small changes to the structure relative to the initial crystal structure. To achieve this, we adapted a Rosetta Script from Pollizi *et. al.* (2017)^33^, used for the design of a photoactive porphyrin-binding tetrahelical bundle. More aggressive backbone sampling methods such as the FastDesign mover^30^ or alternating rounds of sequence design (FixBB) and FastRelax were also tested (Supp. Info.), however although these methods converged to a lower overall Rosetta energy score, molecular dynamics simulations of the apo-m4D2 design suggested that these approaches caused significant disruption to the overall structure.

Conversely, to facilitate the creation of nanoscale protein wires for long range electron transfer, we also designed an extended 4D2 variant, e4D2, capable of binding four heme B’s in a linear array. This required duplication of the diheme 4D2 unit, extending the protein along its helical axis. To ensure appropriate orientation of the heme-ligating histidine side chains into the core of the protein, we separated equivalent histidine residues by 21 residues to fit approximately to three cycles of the helical heptad repeat observed in coiled-coil structures. We built each helix by repeating the sequence of the 25-residue 4D2 helix, removing two residues from each sequence at the junction between repeats, ensuring the correct histidine orientation. This resulted in the 46-residue e4D2 helix, for which we constructed a model using the ISAMBARD^34^ design package. We extracted coiled-coiled parameters from the 4D2 crystal structure and used them to construct the extended helical structure. Finally, we selected the same set of loops from the initial 4D2 design to link the e4D2 helices, and then refined and minimized the model using Rosetta.

### Heme Binding, Helicity and Redox Potentiometry of m4D2 and e4D2

After cloning the mono- and tetraheme m4D2 and e4D2 into the same expression vector as 4D2, we cytoplasmically expressed the proteins in *E. coli* using the same method as for 4D2. Like 4D2, m4D2 binds heme B *in vivo* and retains it through purification, resulting in a ferric UV/visible spectrum indistinguishable from 4D2 (Fig. 5A) and similarly high heme binding affinity (K_D_ <3 nM). In contrast, e4D2 does not bind a significant quantity of heme B *in vivo* under cytoplasmic expression, and we instead primarily purify apoprotein; however, apo-e4D2 readily binds exogenous heme B *in vitro*, exhibiting a similar ferric UV/visible spectrum to both 4D2 and m4D2. Analysis of heme-loaded e4D2 by size exclusion chromatography revealed the presence of aggregated, heme-containing protein, but also a significant quantity of heme-loaded e4D2 eluting at a volume corresponding well to that of a monomeric 25 kDa protein. We found that the yield of monomeric, heme-loaded e4D2 can be improved by adding heme at 37°C, with marked suppression of aggregated or misfolded material relative to additions at 4 and 25°C (Figure 6F). Once separated, this monomeric, heme-bound e4D2 remains stable for several weeks (at 4°C), and further size exclusion chromatography indicated only monomeric e4D2 was present. Given the tendency of purified e4D2 to produce misfolded protein on the addition of heme B, quantification of the heme binding affinity is challenging; however, competition assays using apo horse heart myoglobin^35^ indicate that binding is tight and likely in the nanomolar range (Supplementary Fig. 2). To confirm the heme-binding stoichiometry of m4D2 and e4D2, we performed non-denaturing mass spectrometry on the two designs, revealing the intended 1:1 and 4:1 heme-bound complexes of m4D2 and e4D2 respectively.

**Figure 5:**
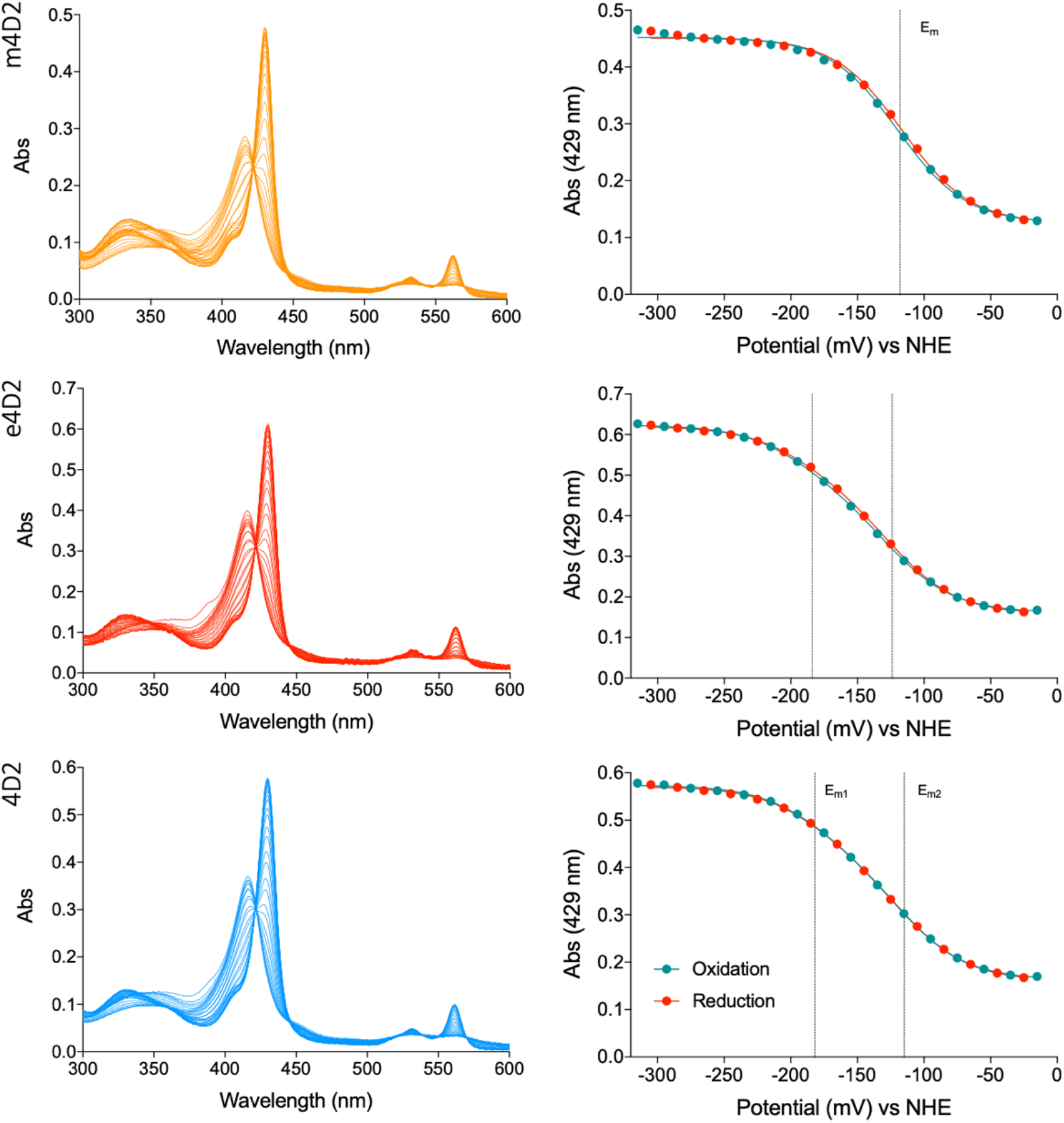
Both the single (m4D2) and four heme (e4D2) variants bind the cofactor, resulting in indistinguishable absorbance spectra compared to the original protein with an oxidized Soret peak at 416nm and reduced peak at 429nm. The overall midpoint of e4D2 is equivalent to that of 4D2 (approximately −150 mV) suggesting equivalent heme environments within the extended, duplicated structure, whereas the midpoint potential of m4D2 (−118 mV) is significantly positively shifted, stabilizing the reduced state of the cofactor. The left panels demonstrate representative UV/Visible spectra of the redox titrations, whilst an average of three repeats are depicted on the right.

**Figure 6:**
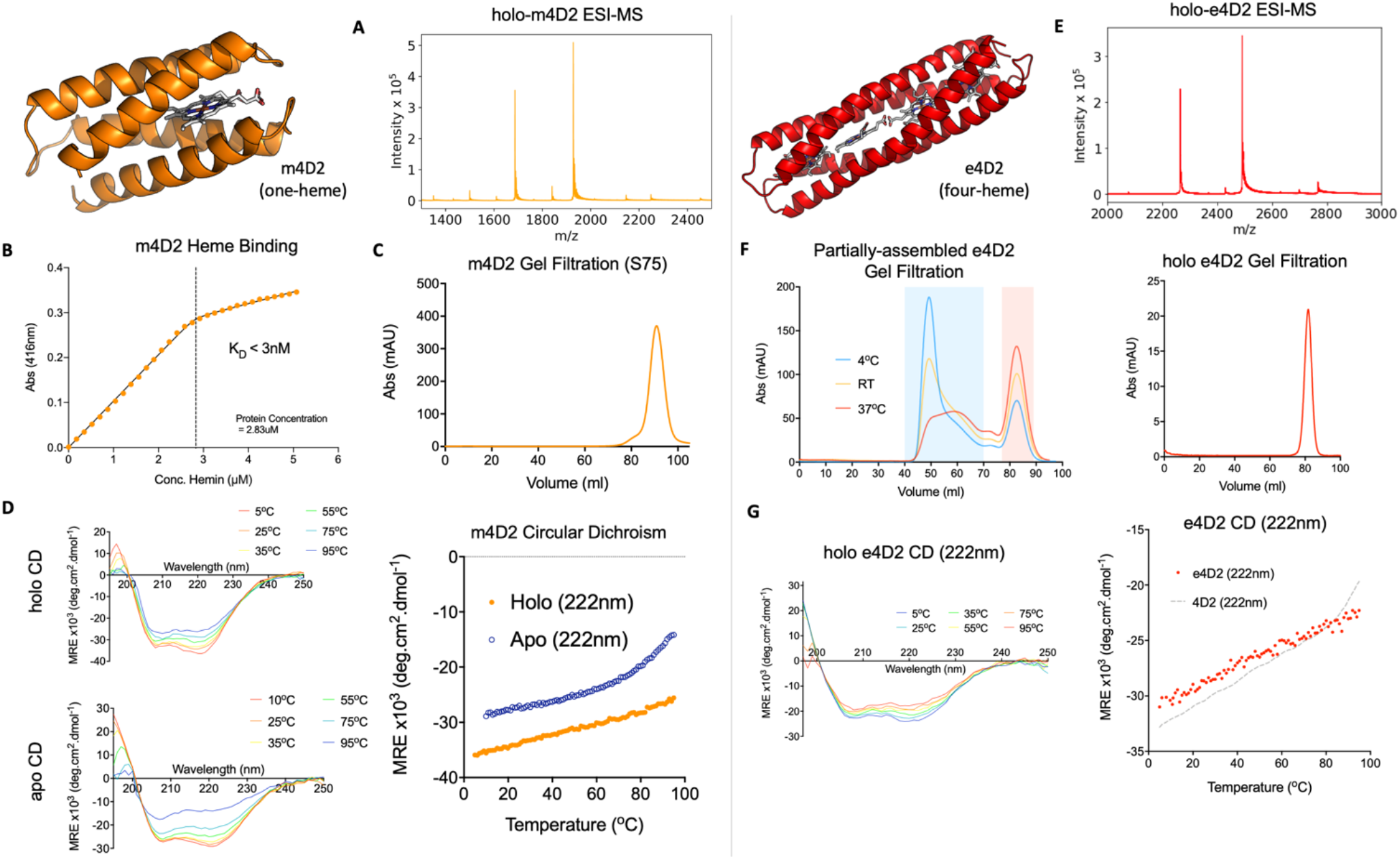
Left panel – assembly of the single heme binding m4D2 variant was confirmed by ESI-MS (A), m/z peaks of 1500.01 (Z=9), 1687.39 (Z=8) and 1928.30 (Z=9) correspond to the total molecular weight of 13,491.09 Da. Tight heme binding and 1:1 stoichiometry was demonstrated by heme titration (B), and homogeneity of the assembly by gel filtration (C). The CD spectra of m4D2 demonstrates a high degree of helicity in both the holo and apo forms, with stability only marginally disrupted above 80°C observed in the apo state. Right panel – similarly, the fully assembly four heme binding e4D2 complex can also be detected by mass spectrometry (Z=9: 2768.17, Z=10: 2492.03, Z=11: 2265.79, total mass 24,909.24 Da). Addition of heme to the purified apo-e4D2 produced a mixture of inhomogeneous aggregates alongside the stable folded complex with the yield improved at increased temperature of heme addition (F), however the monomeric form can be separated by gel filtration and remains highly stable. The secondary structure of e4D2 is very similar to 4D2 compared to the number of residues (G), suggesting successful duplication of the structure.

We then used electrochemistry to provide an insight into the heme environment of these new 4D2 variants (Fig. 5). e4D2 exhibits a broadly similar potentiometric titration to 4D2, with similar overall midpoint potential. While we initially attempted to fit the data to a Nernst function with four single electron redox changes, there were multiple solutions with equal statistical validity and potentials ranging from −120 mV to −195 mV. A possible route to disentangle the potentiometric data would involve selectively modifying or replacing hemes within e4D2 to perturb their individual midpoint potentials, but this is beyond the scope of the present study and future work will focus on achieving this selective modulation of heme potential in a chain of the redox cofactors. For simplicity, we instead used a Nernst function representing two single electron redox events, and observed a slightly smaller separation between midpoint potentials than for 4D2 (−124 and −194 mV for e4D2). This indicates that the hemes are essentially paired in redox potential, with the outer and inner hemes experiencing differing electric field environments as a result of being in close proximity to one or two other hemes respectively. Conversely, m4D2 exhibited a significant positive shift in midpoint potential (Em = −118 mV) when compared to 4D2. This is consistent with the expected electric field effects of placing hemes in close proximity, as previously demonstrated in earlier iterations of the heme-containing maquettes^20,36,37^.

We subsequently employed circular dichroism spectroscopy to probe m4D2 and e4D2 secondary structure and thermal stability. We observed a predictably high degree in helicity for both the monoheme 4D2 and the tetraheme e4D2, and both exhibit excellent thermal stability with no evidence of significant helical unfolding up to 95°C. In contrast to 4D2 and e4D2, apo-m4D2 also exhibits a relatively high degree of helicity and good thermal stability, with a cooperative melt transition starting at approximately 60°C. These observations are consistent with our design methodology and that previously implemented by Degrado *et al* ^33^; we designed m4D2 to contain a well folded core where the second heme binding site was removed, while allowing flexibility around the unoccupied, remaining heme site.

### NMR analysis of the single cofactor m4D2 2,4-DMDPIX complex

Unfortunately, we were unable to obtain diffraction-quality crystals of either holo-m4D2 or e4D2. However, NMR spectroscopy demonstrated that the monoheme m4D2, was well-structured, with good peak dispersion in the 2D H^1^N^15^-HSQC spectrum (Figure 7). Given the observation of alternative heme orientations in the crystal structure of 4D2, we reasoned that substitution of heme B for a symmetrical variant would eliminate such binding heterogeneity and improve NMR signal dispersion. We therefore selected the symmetric heme derivative, iron 2,4-dimethyl-deuteroporphyrin (DMDPIX) for incorporation^38^, as it contains methyl groups in all 1-4 porphyrin positions. DMDPIX binds to m4D2 with slightly lower affinity than heme B (25 nM vs 3 nM respectively), most likely due to the removal of hydrophobic interactions between binding site residues and the vinyl groups of heme B. With DMDPIX bound to double isotopically labelled m4D2 (^13^C^15^N), we were able to obtain 3D NMR spectra that enabled the assignment of >90% of backbone and a large proportion of side chain atoms (Supplementary Table 2). While this data indicates that m4D2 adopts a singular native-like structure, we were unable to obtain sufficient long-range NOE interactions to enable full structural assignment, due to the paramagnetic influence of the symmetric heme. Given these data, we obtained a 2D H^1^N^15^-HSQC spectrum of 4D2 with DMDPIX bound and observed a similar improvement in peak dispersion relative to heme B-bound 4D2, further highlighting the issue of heme binding site heterogeneity. We also obtained 2D NMR spectra of apo-m4D2, demonstrating reasonable peak dispersion which further supports evidence of a well-packed core and relatively flexible/disordered heme binding site when vacant.

**Figure 7:**
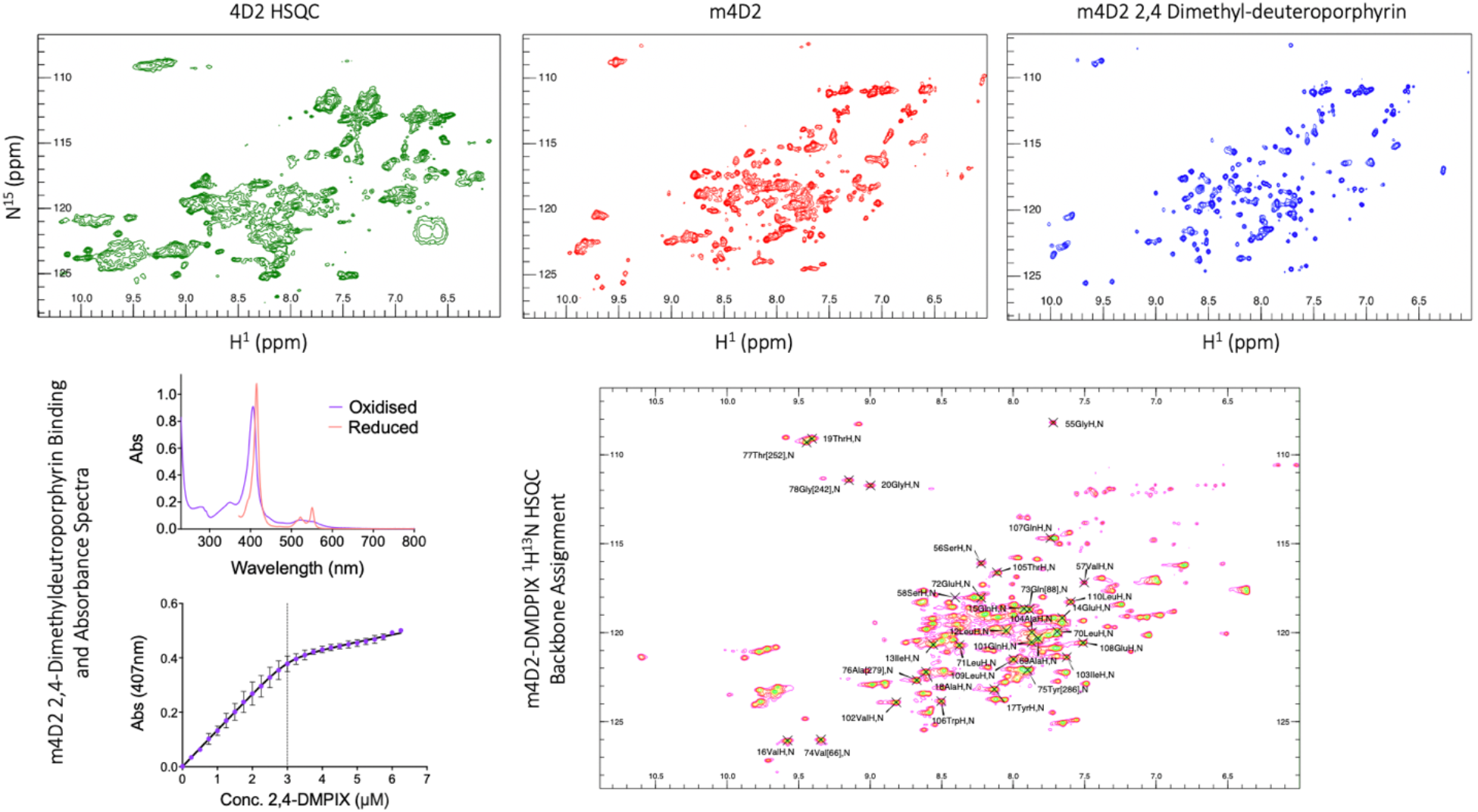
The H^1^N^15^-HSQC spectrum of m4D2 is significantly more dispersed than the two heme original design 4D2, suggesting the single heme protein is also well-structured and further simplified by removal of one of the porphyrins. Peak dispersion was further improved by incorporation of the symmetric heme derivative 2,4-dimethyl deuteroporphyrin, suggesting that heme binding in two asymmetric orientations resulted in broadening of nearby protein signals. The assembly of this cofactor has a slightly blue-shifted absorbance spectra (Soret peaks at 406nm and 415nm in the oxidized and reduced forms) and binds slightly less tightly than heme (K_D_ 25 nM vs 3 nM) however biophysical properties of the complex are otherwise similar. This well resolved spectra alongside 3D NMR experiments enabled comprehensive assignment of peaks (>90%), prediction of secondary structure and torsion angles which correspond to the intended design, and a limited number of NOE based restraints.

### Structural insights into the 25kDa e4D2 by cryoEM

We reasoned that the four heme irons in the tetraheme e4D2 might provide sufficient electron density to observe the protein using electron microscopy. To this end, we initially screened heme-bound e4D2 by negative stain transmission electron microscopy (TEM) and demonstrated evidence of the protein’s extended linear helical structure, with many particles fitting well to the approximate dimensions of 7.5 nm × 2 nm described by the constructed model. Given these TEM results, we then acquired cryo-electron micrographs of tetraheme e4D2. Initially, we collected a dataset of approximately 2000 high quality electron micrographs after optimizing grids for thin ice conditions. Whilst the protein is challenging to identify from cryoEM micrographs, particularly using auto-picking software, we obtained 2D class averages or particles which fit very well to the designed protein specification (Figure 8). These averages show remarkable detail for such a small protein, demonstrating four distinct ‘segments’ along the helical bundle which correspond to the positions of each heme B in addition to depicting four distinct helical structures from a top-view of the molecule through the helical axis. This provides the most substantial evidence that the molecule folds to the designed structure, forming an extended helical bundle with the four tightly packed heme groups positioned in the core of the protein. Preliminary 3D reconstructions of the model demonstrate the overall topology of the helical bundle at low resolution, and aid the pinpointing of the four heme iron atoms. These can be identified as four sites of strong electron density in the refined 3D model, with inter-heme distances that correlate well to the inter-heme distance observed in the 4D2 crystal structure and the molecular dynamics simulations of the e4D2 model.

**Figure 8:**
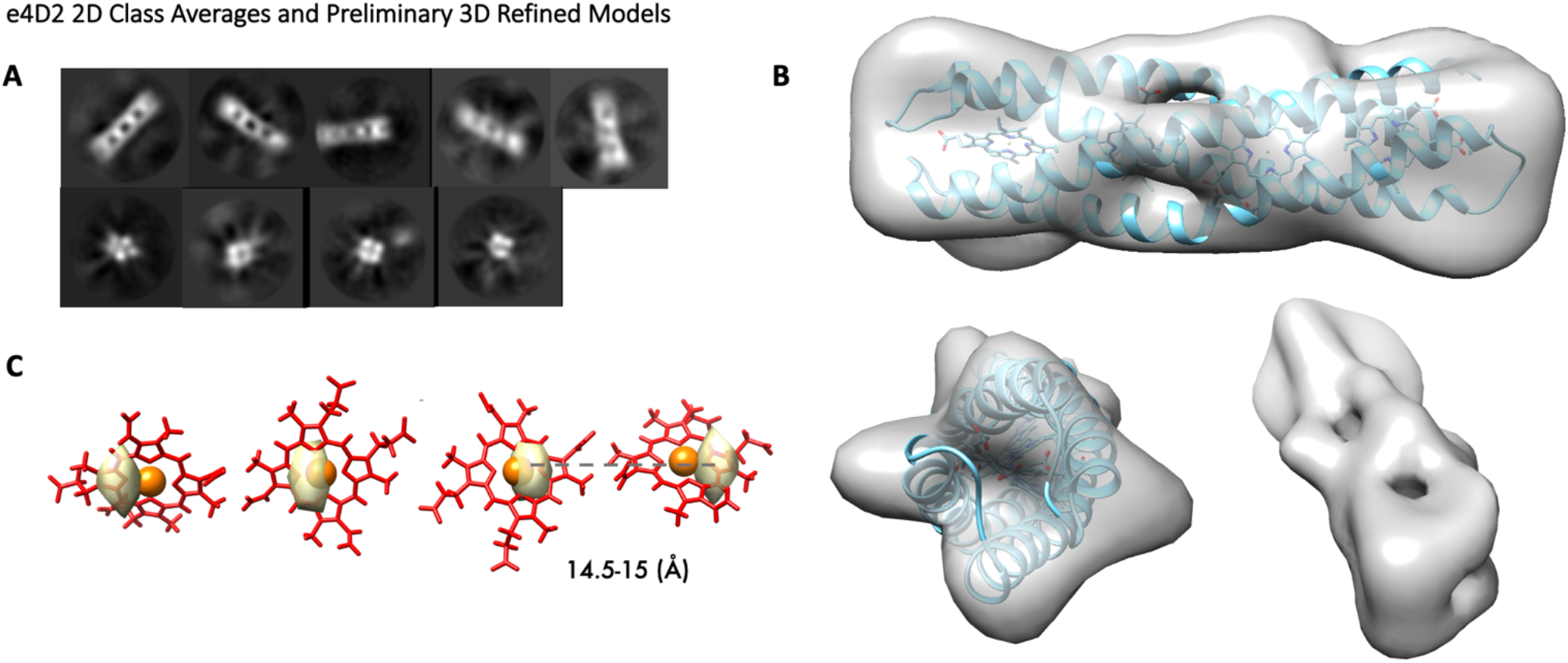
(A) Representative 2D class averages (>500 particles per class) from e4D2 cryoEM micrographs highlighted the four cofactor binding sites along the length of protein (above), in addition to the arrangement of the four helix bundle (below). Initial 3D reconstructions fit closely to the dimensions of the designed protein, and the potential position of the iron atoms can be identified at high contrast (C), suggesting slightly helices which are slightly less coiled than initially modelled resulting in a longer protein and greater separation between the heme groups. This distance matches closely to the equivalent iron-iron distance observed in the crystal structure of 4D2 at 14.7 Å.

### Redox engineering of m4D2

The ability to precisely specify and modulate the midpoint potentials of redox sites and cofactors within proteins would be an exceptionally valuable tool for protein engineers, and *de novo* designed bundles such as the D2 maquettes provide an invaluable framework for developing such a methodology. We therefore selected the monoheme m4D2 for heme redox potential modulation, as it is a simple scaffold with potential for manipulating the electrostatic environment of the heme. To enable the prediction of heme redox potentials in any planned m4D2 variants, we used Monte-Carlo continuum electrostatics (MCCE) calculations^16^ on static models of our proteins, the output providing relative shifts in potential between m4D2 and any altered variants. Furthermore, we initially attempted to both raise and lower the redox potential of the heme, thus establishing this workflow as a means for redox potential prediction and engineering in these proteins.

Previous work with natural peroxidases^39,40^ has established that aspartate residues local to the proximal heme-ligating histidine play key roles in modulating the imidazolate character of the histidine side chain, priming the heme for catalysis by maintenance of the correct histidine tautomeric state. This increase in local electronegativity results in a negative shift of the heme midpoint potential. For m4D2, we selected the two threonine residues involved in ‘keystone’ hydrogen bonding interactions with the proximal histidines for mutation to aspartate, and we constructed both single and double mutant variants (T76D & T18D/T76D). The MCCE calculations predicted shifts in E_m_ of −27 mV for the single aspartate mutant, and −52 mV for the double aspartate mutant (Fig. 9). When we expressed and characterised these designs experimentally, shifts of −27 mV and −55 mV respectively were observed at pH 8.6, closely in agreement with the values predicted by the MCCE calculations.

**Figure 9:**
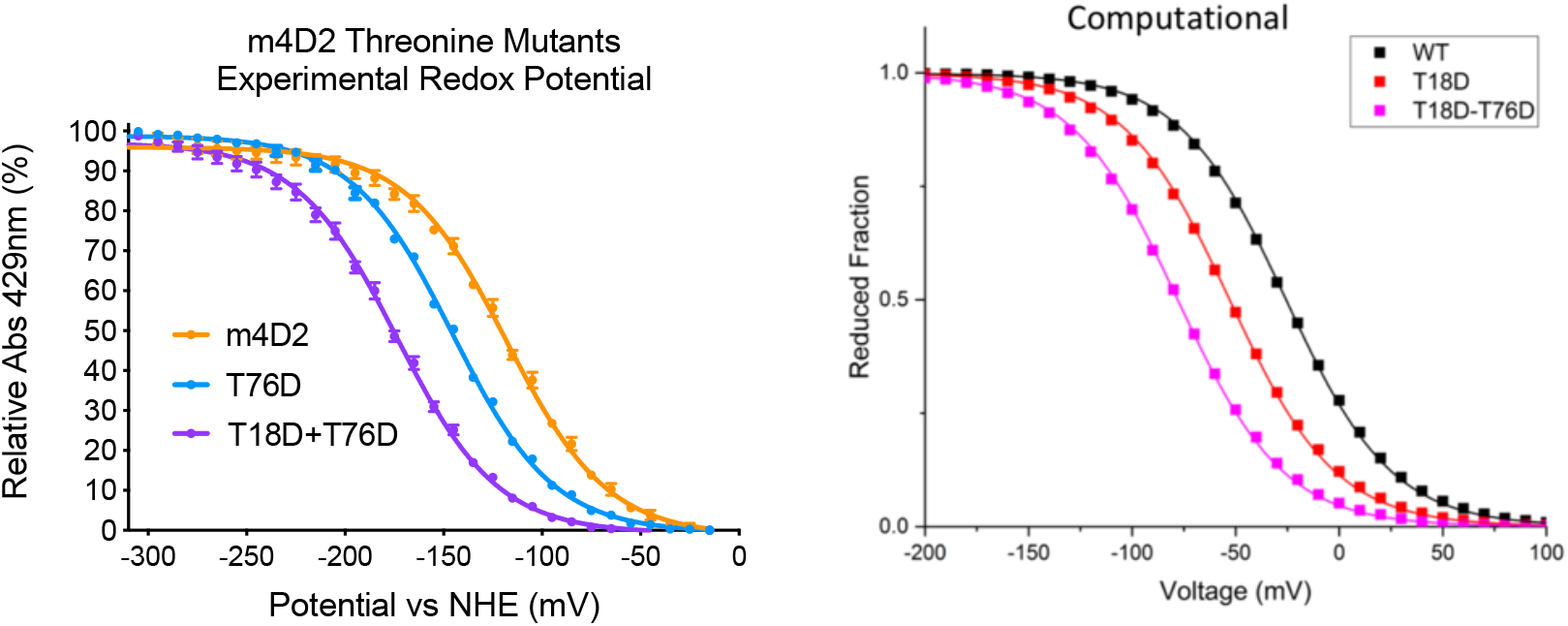
The midpoint potential of the single (T76D) and double (T18D/T76D) threonine mutants of m4D2 displayed midpoints of –145 mV and –173 mV respectively. These negative shifts in midpoint potential of –27 mV for the single mutant and –55 mV for the double mutant were closely aligned to the shifts predicted using MCCE calculations of –27 mV for m4D2-T76D and –52 mV for m4D2-T18D/T76D. The left panel shows the reduction curves for an average of three experimental repeats fitted with a single-electron Nernst fit, while the right panel shows the reduction curves predicted computationally using the MCCE method.

Future work will focus on more wholesale remodeling of m4D2 and the multi-heme 4D2 and e4D2 proteins to engineer greater changes in the electrostatic environments of these maquettes. These calculations can also be incorporated into a pipeline for directed modulation of the behavior of these simple proteins and help us to elucidate how the local protein environment impacts the behavior of the hemes within them. Significantly, we may be able to engineer directional midpoint gradients through multi-heme maquettes, with potential applications in electron transport through nanoscale protein-based wires.

## Discussion and Conclusions

We have attempted to address the deficiencies in high resolution design of *de novo* redox proteins by creating a framework of single and multi-heme proteins with apparently well-defined structural attributes. These offer much potential in exploring fundamental biophysical properties and functions of cofactors and natural oxidoreductases, such as electron transfer and possibly catalysis, while providing stable, robust scaffolds that could play a role in the modular assembly of functional nanoscale objects for energy capture and generation. A key feature in design is predictability; without this, atomistically precise protein design is not possible, and gaining complete and fine control of protein and cofactor properties relies on a potentially exhaustive, iterative design process with no guarantee of success. To this end, the structure of 4D2 matches well that of the predicted structure of D2^15^, which is a testament to the computational methods originally used in the design of the self-assembling peptide. D2 was originally designed by parametrization of the transmembrane cytochrome *b* of the cytochrome bc1 complex, and the crystal structure of 4D2 reveals an excellent reproduction of diheme orientation and spacing in a robust, mutable and soluble protein.

This scaffold also provides opportunities in the further design and engineering of such proteins. Removal of a heme binding site alongside compensatory redesign of the pocket to facilitate improved core packing, resulted in a monoheme protein that appears well-folded and retains attractive features from the original design. Since 4D2 was essentially cleaved into two separate halves for the production of m4D2, it is possible that such proteins can be designed in a modular process, where parts with differing biophysical attributes (e.g. redox potentials) can be combined to dictate, for example, the direction of electron transfer along the helical axis, or to create multi-cofactor proteins with specificity in each individual binding site. This is especially applicable to the larger tetraheme e4D2, where directionality could potentially be imparted onto the hemes to dictate electron transfer.

By expanding the protein in this way, we were also able to push the current boundary of cryo-EM^41,42,43^, principally facilitated by the electron-rich, heme-saturated core. While the data falls short of atomistic assignment, it does offer robust evidence of the global protein structure, which fits the dimensions of the small, but extended, helical bundle and the hemes located in the core. The assembly of the protein requires additional purification after heme loading, but production is reliable and the design highly stable, offering a scaffold for the design of properties such as directional electron transport through the 8 nm length of the protein. Our design and modification of these derivatives demonstrates the adaptability of the initial 4D2 protein, offering a platform for structure-based design and modification from crystal structure to function.

## Acknowledgements and funding

This work was supported at the University of Bristol by the Biological and Biotechnological Sciences Research Council ([BBI014063/1], [BB/R016445/1], [BB/M025624/1], [BB/M009122/1] & [BB/T008741/1], the latter two providing a studentship for G.H.H.) and the SynBioCDT (EPSRC and BBSRC Centre for Doctoral Training in Synthetic Biology Grant EP/L016494/1) for studentships for C.E.M.N. and B.H. The Authors would also like to thank Dr Peter Wilson in the School of Biochemistry Biosuite at the University of Bristol for access to biophysical equipment and Dr Ufuk Borucu for assistance with Cryo-EM data acquisition. The Authors declare no conflict of interest.

## Methods

### Molecular Biology

Unless indicated, designs were expressed in the cytoplasm within a pET151 vector (Thermo Fisher) with an N-terminal 6xHis tag and TEV cleavage sequence. Periplasmic expression was utilized to boost non-covalent heme incorporation, facilitated by expression in a modified pMal-p4x vector (NEB), pSHT^18^, including an additional N-terminal translocon recognition sequence. Gene sequences were synthesized by Eurofins Genomics, and cloned by a Site-Directed Ligase Independent Cloning^44^ method, using Q5 polymerase (NEB) to amplify gene and vector sequences prior to hybridization of overhanging fragments. Cloned vectors were transformed into Stellar cells (Takara) prior to plasmid purification at a concentration of 30-100 ng/μL using an NEB miniprep kit.

### Protein Expression and Purification

All proteins were expressed in the NEB T7 Express *E. coli* strain. Transformed colonies were cultured overnight in 100 ml LB with carbenicillin at a concentration of 50 μg/ml, and 20 ml of culture transferred to 1L of LB which was grown at 37°C to an OD_600nm_ of 0.6 prior to induction by addition of 1 mM IPTG. Proteins were expressed for four hours at 37°C, after which cells were harvested by centrifugation at 4000xg for 25 minutes and resuspended in lysis buffer (50 mM Sodium phosphate, 300 mM Sodium chloride, 20 mM imidazole, pH 8).

Isotopically labelled samples for NMR analysis were expressed in minimal media using a method adapted from Rupasinghe *et al.* (2007)^45^. Proteins were labelled by addition of ^15^NH_4_Cl (Sigma) alone or in addition to ^13^C-Glucose (Cambridge Isotopes) for 3D spectra acquisition. Cells were grown to an OD_600nm_ of 0.6 as described above prior to centrifugation and washing in an M9 salts solution and then resuspended in 250ml minimal media with ^15^NH_4_Cl and 4 g/L ^13^C-Glucose/Glucose with M9 salts and trace metals. The full composition of the minimal media solution was as follows: 18 mM ^15^NH_4_Cl, 4g/L glucose, 34 mM Na_2_HPO_4_, 22 mM KH_2_PO_4_, 86 mM NaCl, 2 mM MgCl_2_, 50 μM FeCl_3_, 20 μM CaCl_2_, 10 μM MnCl_2_,10 μM ZnSO_4_, 2 μM CuCl_2_, 2 μM CoCl_2_, 2 μM NiCl_2_, 2 μM H_3_BO_3_. Cultures were incubated at 28°C for one hour to aid cell recovery before the addition of IPTG, and protein expressed for 16-18 hours at 28°C.

Resuspended cell-pellets were lysed by four 20-second sonication steps on ice, and cell debris removed by centrifugation at 18,000 xg. Designs were initially purified from lysate by nickel affinity chromatography (HisTrap column GE), eluting at an imidazole concentration up to 250 mM. The proteins were immediately dialyzed overnight (SnakeSkin 3.5 kDa MWCO) into a low salt buffer optimised for TEV cleavage (50 mM Tris, 0.5 mM EDTA, pH 8), followed by cleavage of the N-terminal tags by addition of approximately 200 μg of TEV protease per 3 L of culture media and the reducing agent TCEP to a concentration of 1 mM, under anaerobic conditions. Cleaved designs were further purified by reverse nickel chromatography, followed by size exclusion chromatography (GE HiLoad 16/600 Superdex 75pg) under final buffer conditions of 20mM CHES and 50mM Potassium chloride, pH 8.6. Purified protein samples were concentrated using centrifugal concentrators with a molecular weight cut-off of 3 kDa (Vivaspin /Amicon).

If necessary, exogenous heme was added *in vitro* to fully saturate cofactor binding sites by dropwise addition of hemin dissolved at 1 mg/ml in DMSO to a small stochiometric excess. Excess heme was removed by gel filtration using G25 desalting columns or a further round of size exclusion chromatography. Fully apo samples were prepared by acidic butanone extraction of the heme cofactor followed by dialysis and gel filtration back into buffer.

### Porphyrin Binding Titrations

Cofactor binding titrations were carried out by preparation of up to 1000 μL of 2-3 μM apo protein solution in a quartz cuvette, followed by incremental addition of porphyrin in low volumes of DMSO (0.5-2 μL). Full absorbance spectra between 200-800 nm were measured after each addition and thorough mixing. Binding affinity (K_D_) was calculated by fitting absorbance at a single wavelength (the Soret peak) to the tight binding quadratic equation.

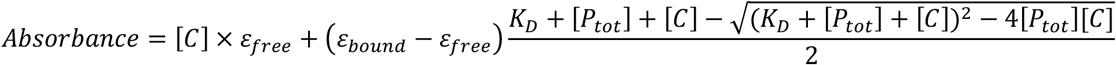

[*C*] = Cofactor Concentration (μM)

*ε*_*free*_ = Unbound Cofactor Extinction Coefficient (μM^−1^cm^−1^)

*ε*_*bound*_ = Bound Cofactor Extinction Coefficient (μM^−1^cm^−1^)

[*P*_*tot*_] = Total Protein Concentration (μM)

### Mass Spectrometry

The molecular weight of heme bound protein complexes was measured using electrospray ionization-mass spectrometry (ESI-MS). Data was acquired using a Waters Xevo G2-XS QTof LC-MS instrument – bypassing the liquid chromatography system by direct injection purified protein samples at a concentration of 50-200 μM prepared in an aqueous 100mM ammonium acetate solution. The sample was injected at a flow rate of 0.25 ml/minute of ammonium acetate, with a capillary voltage of 3.6 kV and cone voltage of 50 V, in positive ion mode. Predicted ion m/z values were calculated by the following equation, assuming the ionization occurred by gaining of a proton, increasing the mass (m) by 1 gmol^−1^ per charge (Z).

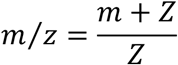

The molecular weight of apo proteins was validated by Matrix-assisted laser desorption/ionization (MALDI) MS, acquired on Bruker UltrafleXtreme equipment. Protein samples (concentration approximately 50 μM) were dissolved 1:10 in a matrix solution of 10mg/ml sinapinic acid in 50:50 water/acetonitrile with 0.1% trifloroacetic acid.

### Redox Potentiometry

Heme redox potentials were measured by OTTLE potentiometry (Optically transparent thin-layer electrochemistry) as described previously^23^. All designed proteins were prepared at a concentration of 50 μM in 10% glycerol, 20mM CHES, 100 mM KCl at a pH of 8.6. A total of six redox mediators were added at the following concentrations: (Phenazine Ethosulfate 20 μM, Indigotrisulfonate 50 μM, Duroquinone 6 μM, 2-Hydroxy-1,4-Napthoquinone 25 μM, Phenazine 20 μM, Anthroquinone-2-sulfonate 20 μM). UV-visible spectra were measured using a Cary 60 spectrophotometer whilst altering the potential via a thin platinum electrode between −225 to −525 mV vs a silver chloride electrode, controlled by a Biologic SP-150 potentiostat. Potentials were adjusted vs the Nernst hydrogen electrode (NHE) by calibration with cytochrome C, resulting in an average adjustment of +220 mV. Midpoint potentials (E_m_) were derived by fitting oxidation and reduction data to the following one or two electron Nernst equations, where mV is the applied potential and Abs is the absorbance at the reduced Soret peak (approximately 429 nm).

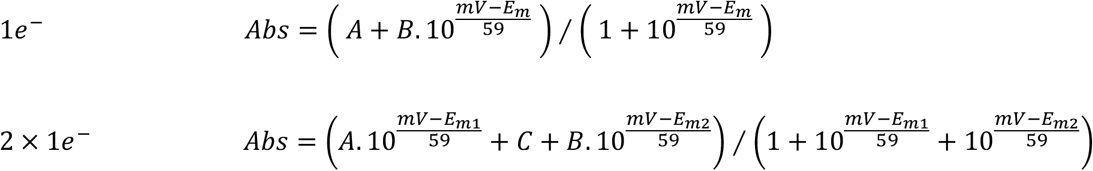

### Circular Dichroism

Circular Dichroism (CD) spectra were recorded using a JASCO J-1500 spectrophotometer. All protein samples were prepared at a concentration of approximately 5-15 μM, and CD spectra baselined against the buffer solution (CHES 20 mM, KCl 100 mM, pH 8.6). CD signal was measured at 222 nm whilst changing temperature between 5-95°C, at a rate of 1°C per minute, to track the overall helicity of designs. Full CD spectra were measured between 190-250 nm at specific temperature points, acquiring eight spectra at each condition and averaging the results. Circular dichroism was reported as mean residual ellipticity (MRE) – converting raw data by the following equation (where n is the number of peptide bonds in a protein sample):

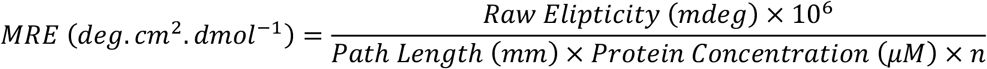

### X-Ray Crystallography

Crystals suitable for data collection were obtained by the sitting drop vapour diffusion method at 22°C using the Morpheus screening conditions (Molecular Dimensions Ltd). Protein samples were concentrated to between 10-20 mg/ml and mixed in a 1:1 ratio with precipitant in 1 μl sitting drops. X-ray diffraction data were collected at beamline I03 (Diamond Light Source). A fluorescence edge scan was measured to determine required wavelengths for phase solving by multi-wavelength anomalous dispersion (MAD), collecting diffraction data at peak, inflection and high remote wavelengths (*1.738 Å, 1.741 Å, 1.698 Å*). Diffraction data were integrated and scaled with XDS and XSCALE using xia2 prior to phasing and model refinement in the ccp4i2^46^ software suite. Experimental phasing by MAD was performed using SHELLX, followed by iterative model refinement using Refmac5 and Coot. 4D2 crystallized in the H3 space group with unit cell dimensions *a=81.25 Å, b=81.25 Å* and *c=59.06 Å*. The refined coordinates have been deposited to the PDB (*ID: 7AH0*) and data collection and refinement statistics are available in Supplementary Table 1.

### NMR

Protein samples were concentrated to between 0.5-2 mM in 10% D_2_O, and NMR spectra were obtained with a 700MHz Bruker Avance III HD instrument (BrisSynBio NMR facility). Spectra were analyzed using Ccpnmr Analysis7 Version 2.4^47^.

### Electron Microscopy

Purified protein samples applied to CF300-Cu carbon coated grids (Electron Microscopy Sciences) were prepared for negative stain transmission electron microscopy, pre-treating grids by glow discharge. Protein concentrations used were between 0.01-0.05 mg/ml, and samples were stained with 2% uranyl acetate. 200 images were obtained at magnification factors between 50,000-90,000x, using a Tecnai 20 electron microscope. Individual particles were picked, and images processed using Scipion software^48^ to produce 2D class averages.

Cryo-electron microscopy micrographs were obtained with a 200kV FEI Talos Arctica (GW4 CryoEM Facility). Quantfoil R 1.2/1.3 grids were prepared with a protein concentration of 2 mg/ml. All analysis was completed using Relion 3.0^49^, including motion correction, CTF estimation, particle picking and 2D and 3D averaging and model building.

### Molecular Dynamics

MD trajectories were calculated in AMBER^50^. The Amber ff14SB forcefield was used to parameterize protein residues, whilst parameters derived for the bis-histidine ligated b-type heme in Cytochrome C oxidase^51^ were used for the cofactors. GPU accelerated production MD was run using the pmemd.cuda application on University of Bristol HPC clusters (Bluecrystal phase 3, 4, BlueGem). Equilibration and molecular dynamics input files and parameters are included in the Supplementary Information. Trajectories were combined and analysed using ccptraj.

### Monte-Carlo Continuum Electrostatics (MCCE)

Models of the m4D2 mutants were generated using PyMOL. Structures were solvated and energy minimisation performed using GROMACS 5.2.1 with the GROMOS 54a7 force field^52^. Restraints were first set to heavy atoms and then Cα carbon atoms, and finally removed entirely. Histidines 36 and 94 were bonded to the heme as part of the redox centre and separated from the other titratable sites, and propionate sites were treated as titratable sites. The continuum electrostatics-based protonation energies were calculated using the MEAD software package, at an ionic strength of 50 mM and a temperature of 300 K. Multiple focused grids were constructed, with spacings of 1, 0.5 and 0.25 angstroms respectively. The dielectric constant for protein was 20 and for solvent 80. Monte Carlo sampling of protonation states was done using the PETIT program. The pH was varied between 8.4 and 8.8 at an interval of 0.2, the potential varied from −250 to 250 mV at an interval of 10 mV, and the temperature was 310 K.

## Supplementary Information

**Supplementary Figure 1:**
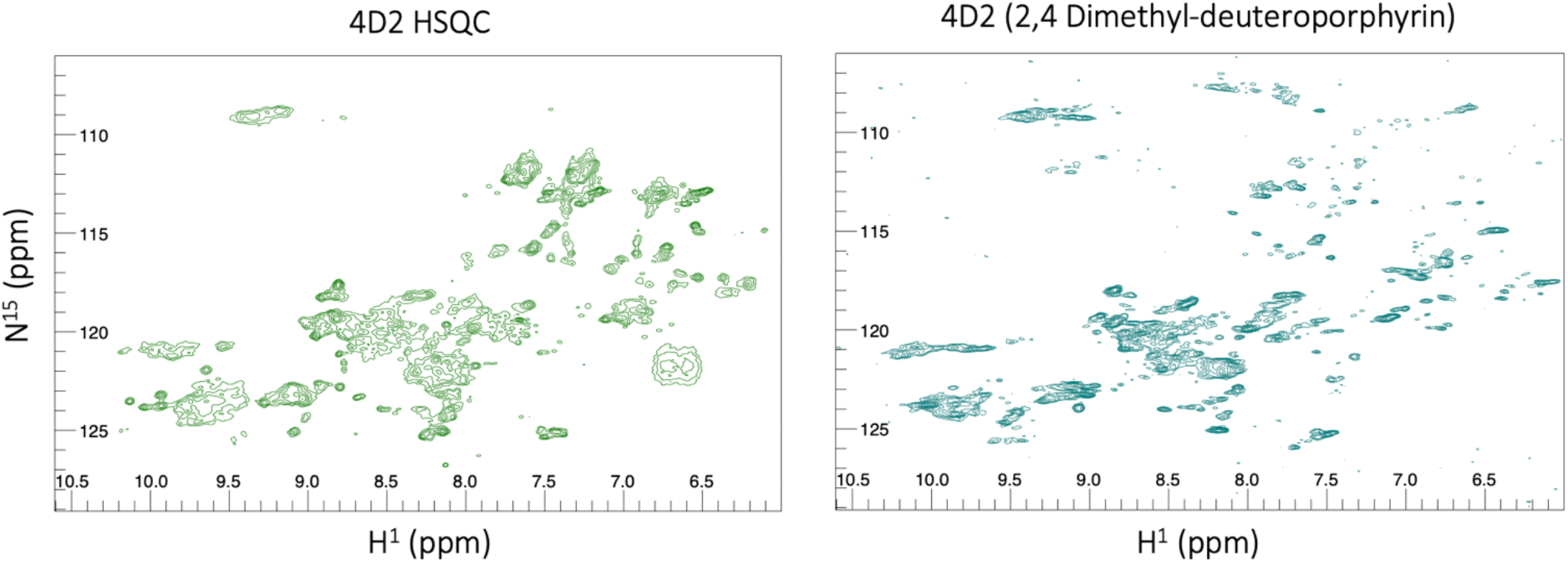
4D2 ^1^H^15^N-HSQC spectrum demonstrates moderate peak dispersion, insufficient for sequence assignment or further structural analysis. Reconstitution with a symmetric porphyrin (right) results in a significant shift, offering further evidence for multiple heme binding states, but does not improve the spectrum quality to the same extent as observed for the monoheme variant m4D2.

**Supplementary Figure 2:**
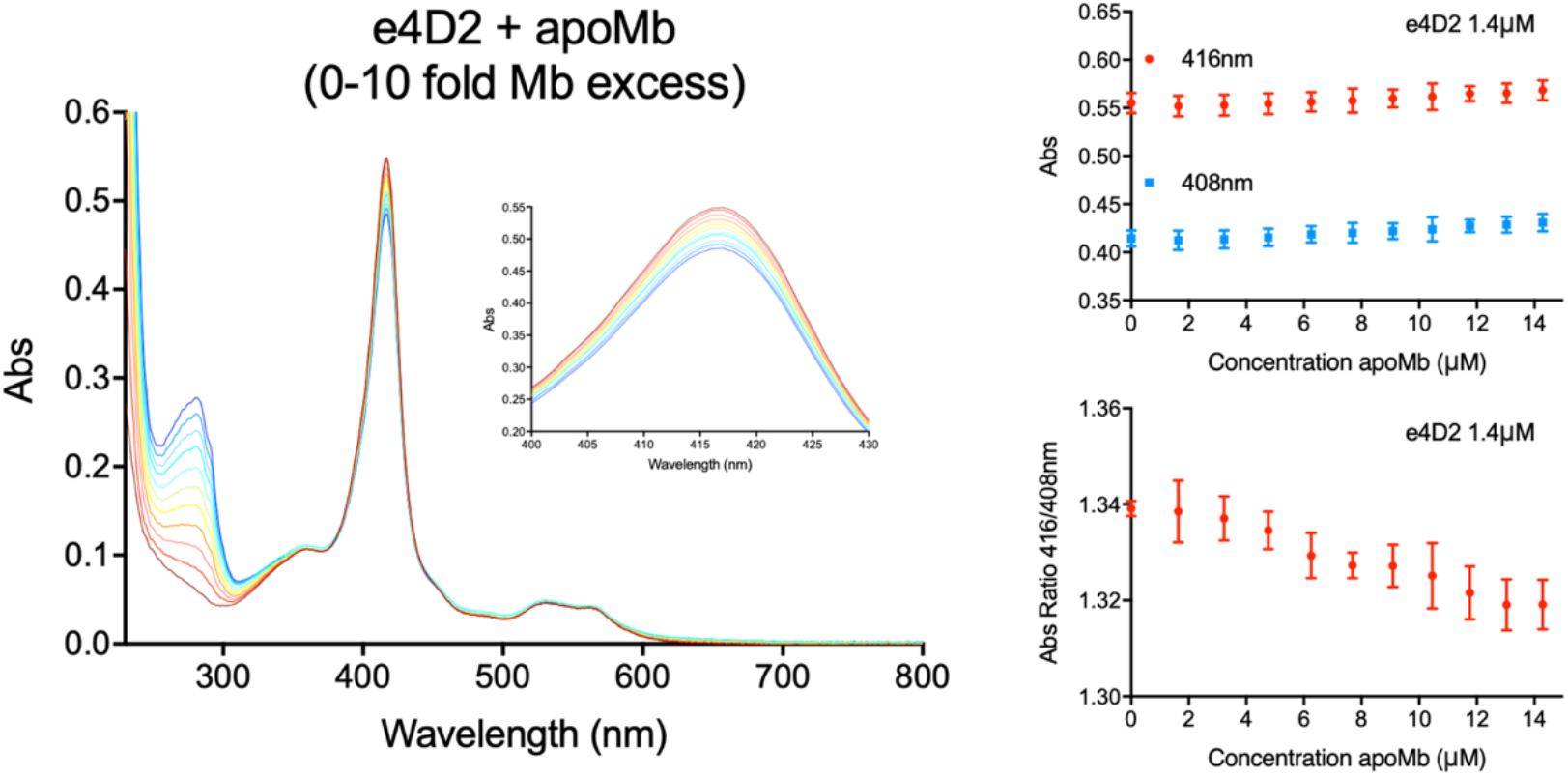
Quantifying heme binding affinity of e4D2 is challenging due to the mixed population of bound states that form upon heme addition. However, negligible transfer of heme to apo-myoglobin (apoMb) is observed after one hour of mixing at an excess of up to 10-fold myoglobin, comparable to heme binding proteins with nanomolar affinity for heme^35^. Complete transfer of heme to myoglobin would result in a 416nm/408nm ratio of 0.77, however only a 1-2% shift of this ratio is detected. This further supports tight binding of heme to e4D2.

**Supplementary Figure 3:**
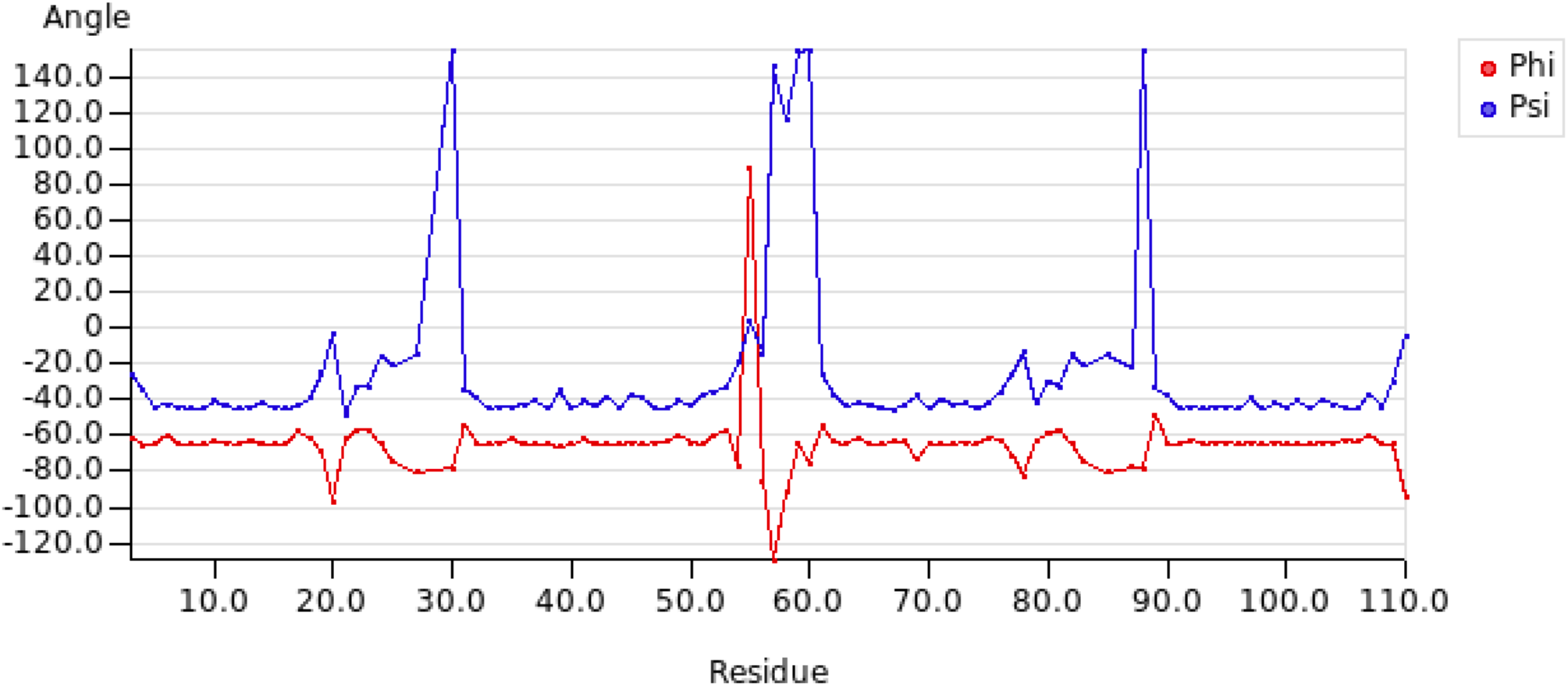
Torsion angle predictions from NMR assignments (calculated using DANGLE in CCPNMR Analysis) indicate the four ordered helical regions connected by three loops. Deviations at residues 20 and 78 are both glycine.

**Supplementary Figure 4:**
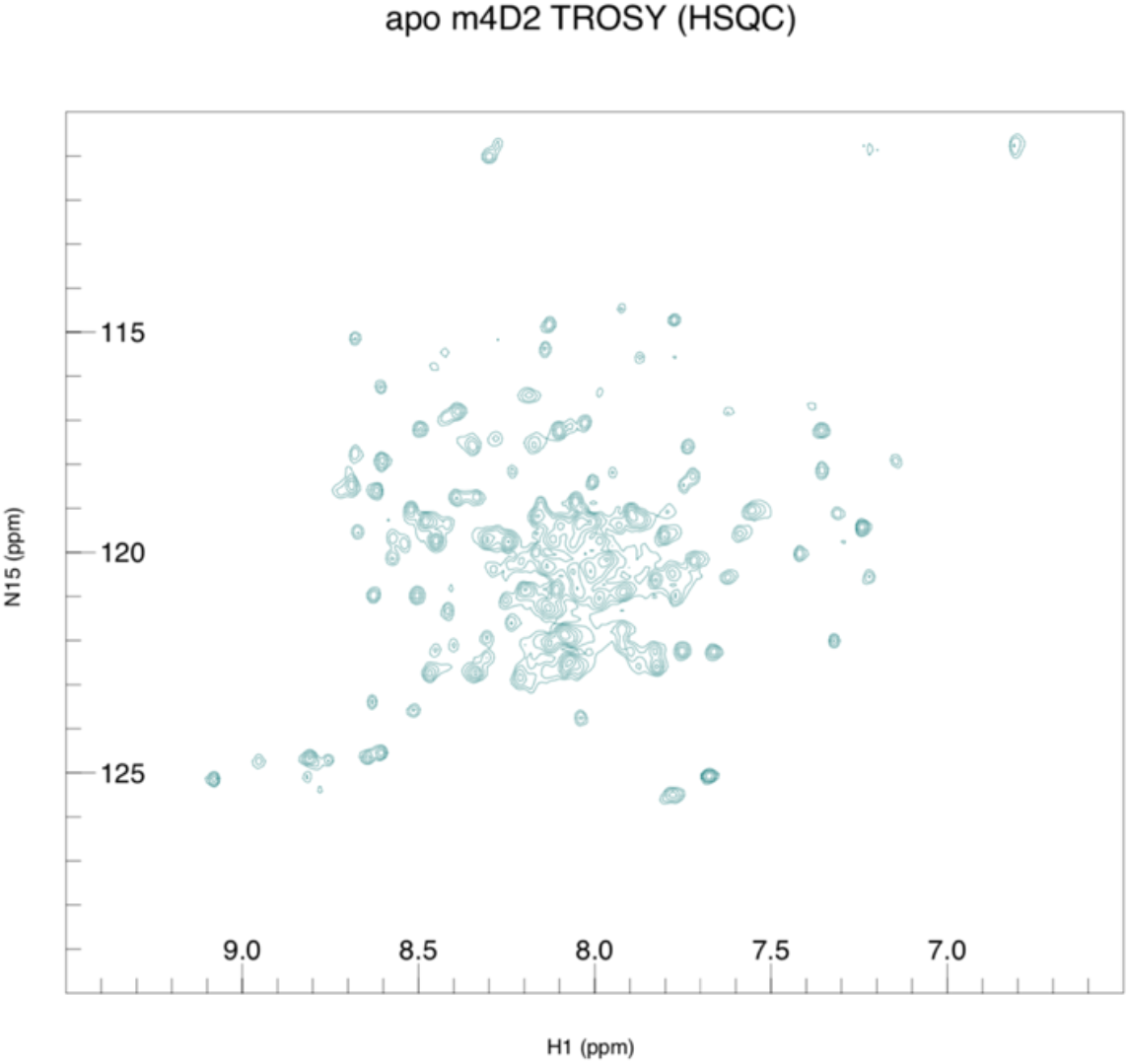
The HSQC (TROSY experiment) for apo m4D2 confirmed that the protein is relatively well structured without bound heme with reasonable peak dispersion, although the repetitive sequence is likely to result in significant overlap of signals

**Supplementary Table 1.**
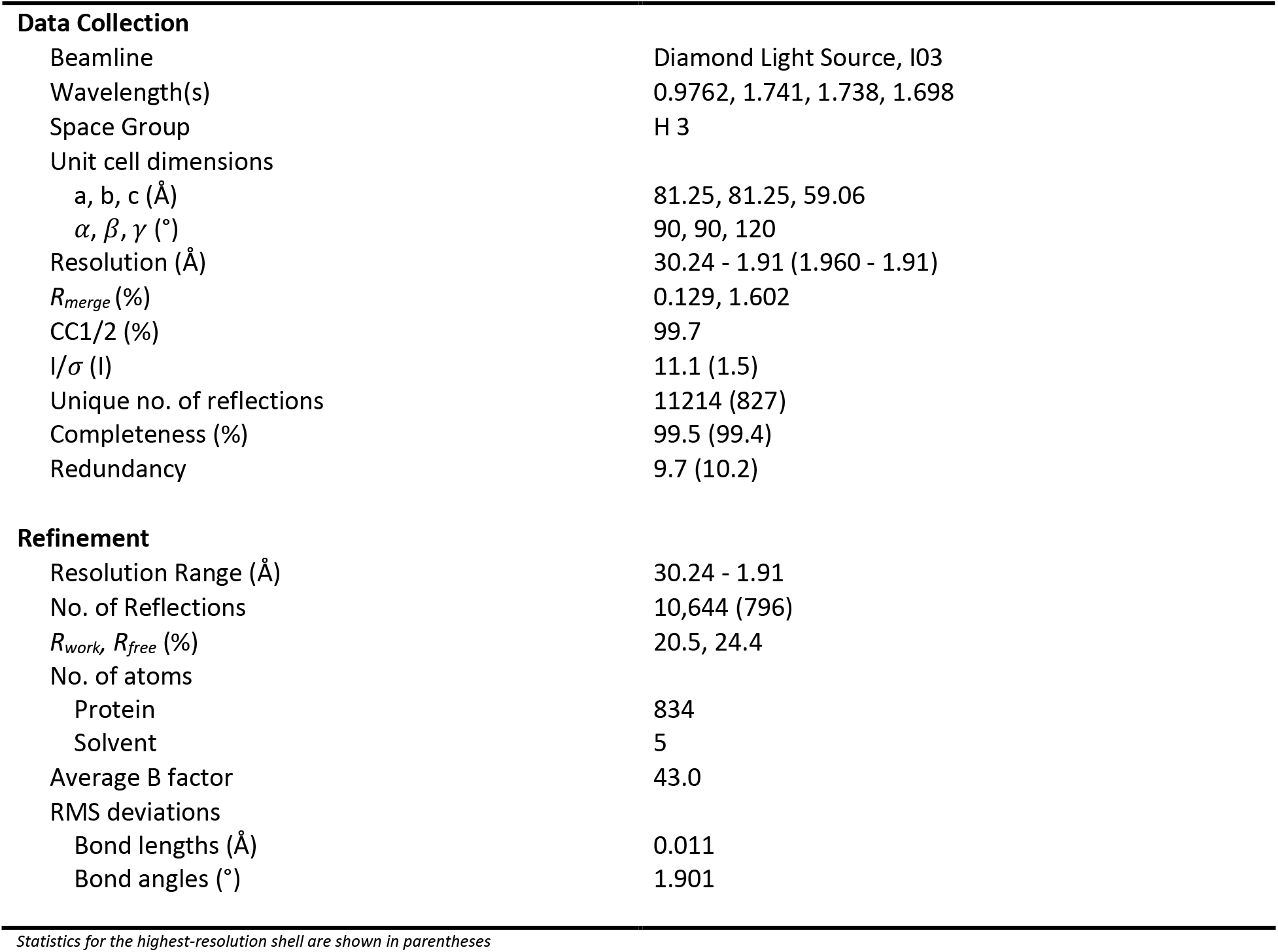
Data collection and structural refinement statistics of 4D2.

**Data Collection**
|  |  |
| --- | --- |
| Beamline | Diamond Light Source, I03 |
| Wavelength(s) | 0.9762, 1.741, 1.738, 1.698 |
| Space Group | H 3 |
| Unit cell dimensions |  |
| a, b, c (Å) | 81.25, 81.25, 59.06 |
| $\alpha$, $\beta$, $\gamma$ (°) | 90, 90, 120 |
| Resolution (Å) | 30.24 - 1.91 (1.960 - 1.91) |
| $R_{\text{merge}}$ (%) | 0.129, 1.602 |
| CC1/2 (%) | 99.7 |
| I/$\sigma$ (I) | 11.1 (1.5) |
| Unique no. of reflections | 11214 (827) |
| Completeness (%) | 99.5 (99.4) |
| Redundancy | 9.7 (10.2) |
| **Refinement** |  |
| Resolution Range (Å) | 30.24 - 1.91 |
| No. of Reflections | 10,644 (796) |
| $R_{\text{work}}$, $R_{\text{free}}$ (%) | 20.5, 24.4 |
| No. of atoms |  |
| Protein | 834 |
| Solvent | 5 |
| Average B factor | 43.0 |
| RMS deviations |  |
| Bond lengths (Å) | 0.011 |
| Bond angles (°) | 1.901 |

*Statistics for the highest-resolution shell are shown in parentheses*

**Supplementary Table 2.**
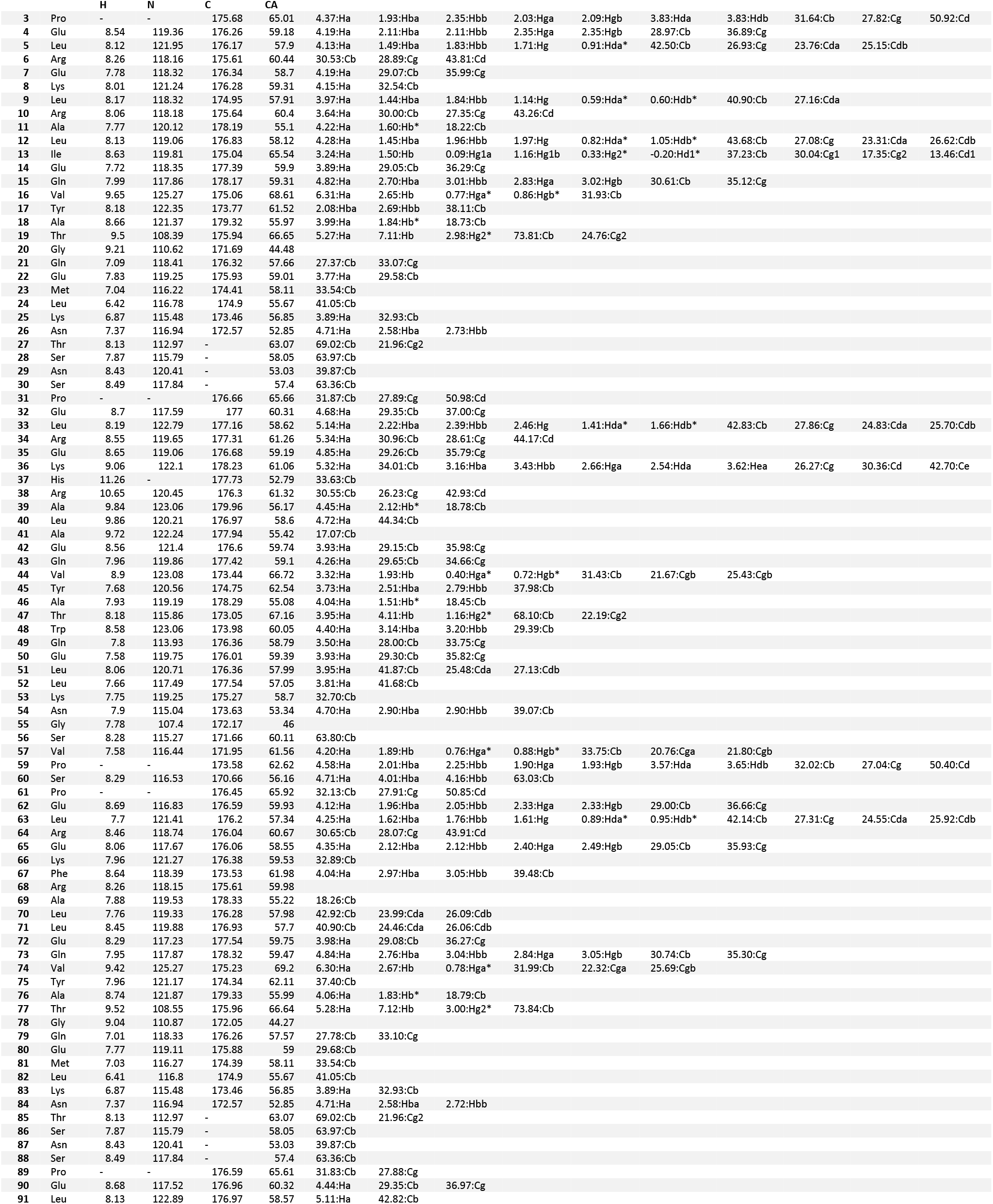
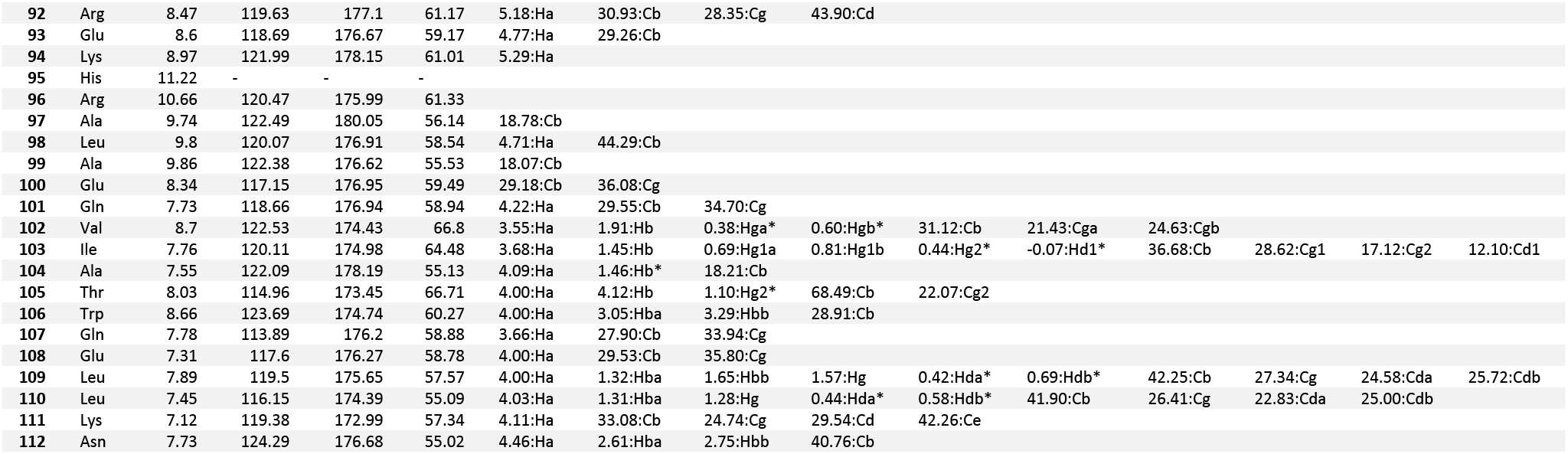
NMR chemical shifts for m4D2 (2,4-Dimethyldeuteroporphyrin)

## m4D2 Design Rosetta Scripts

### FastDesign.xml

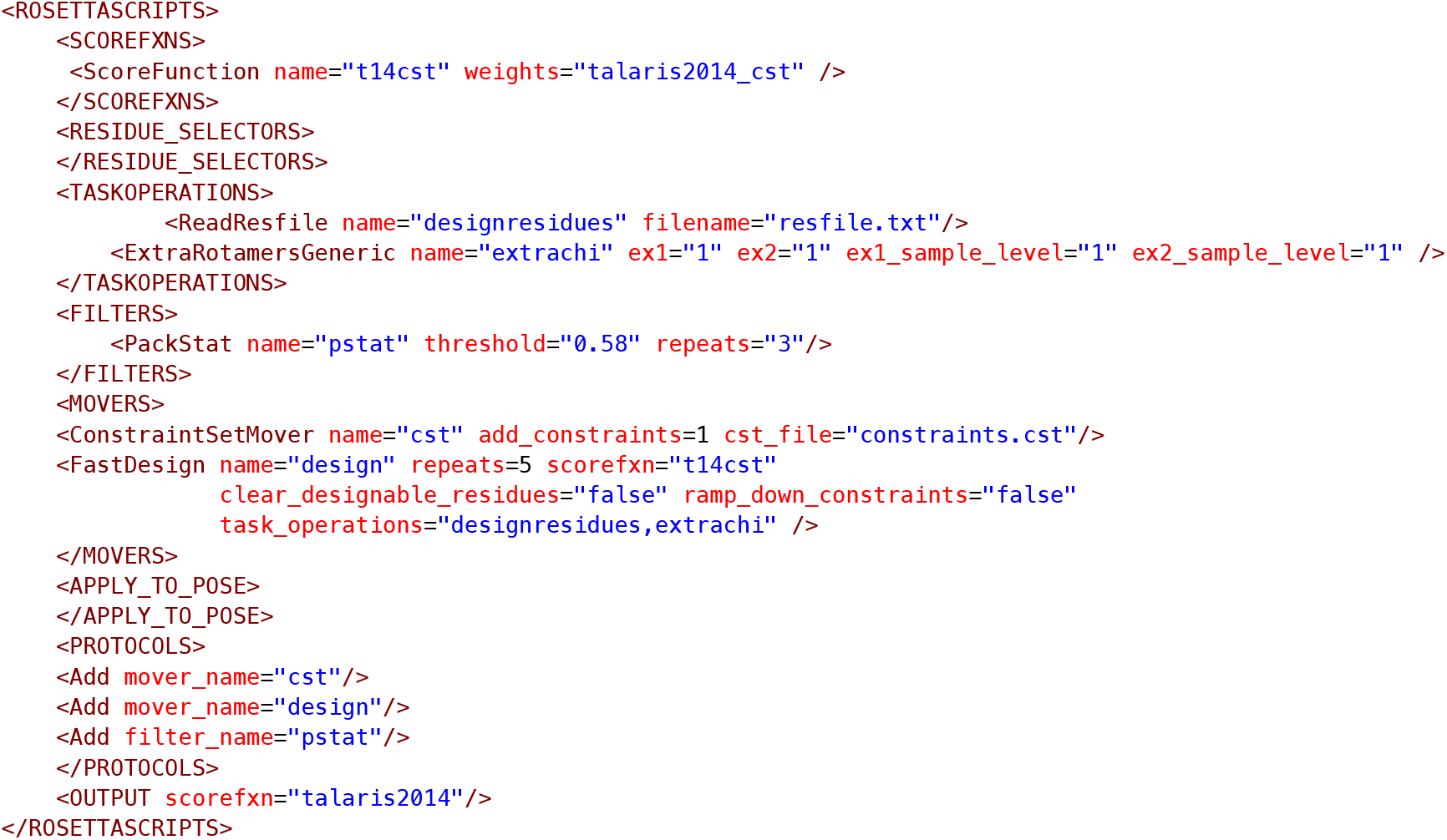

### Relax-Repack.xml

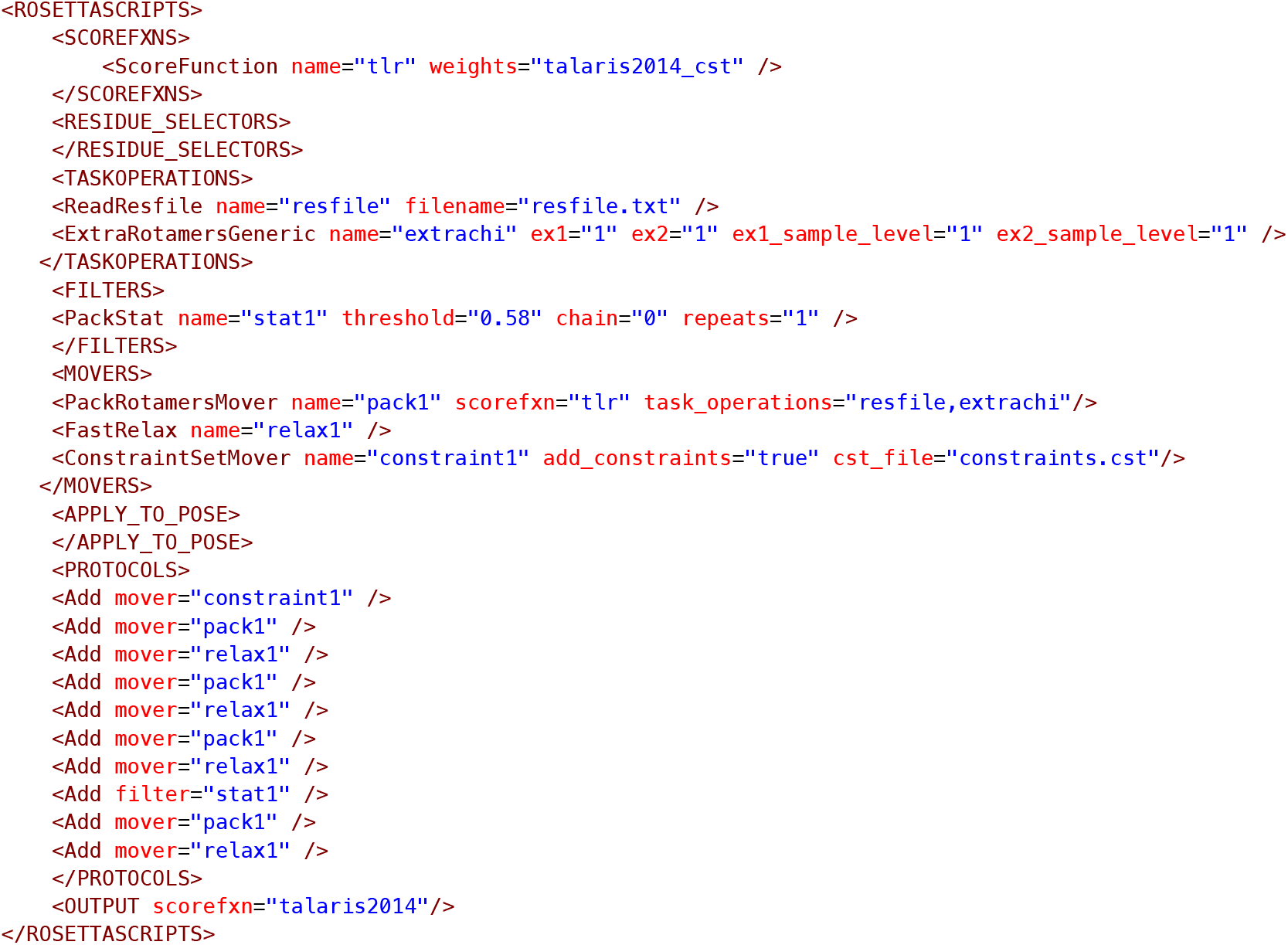

### Backrub.xml^33^

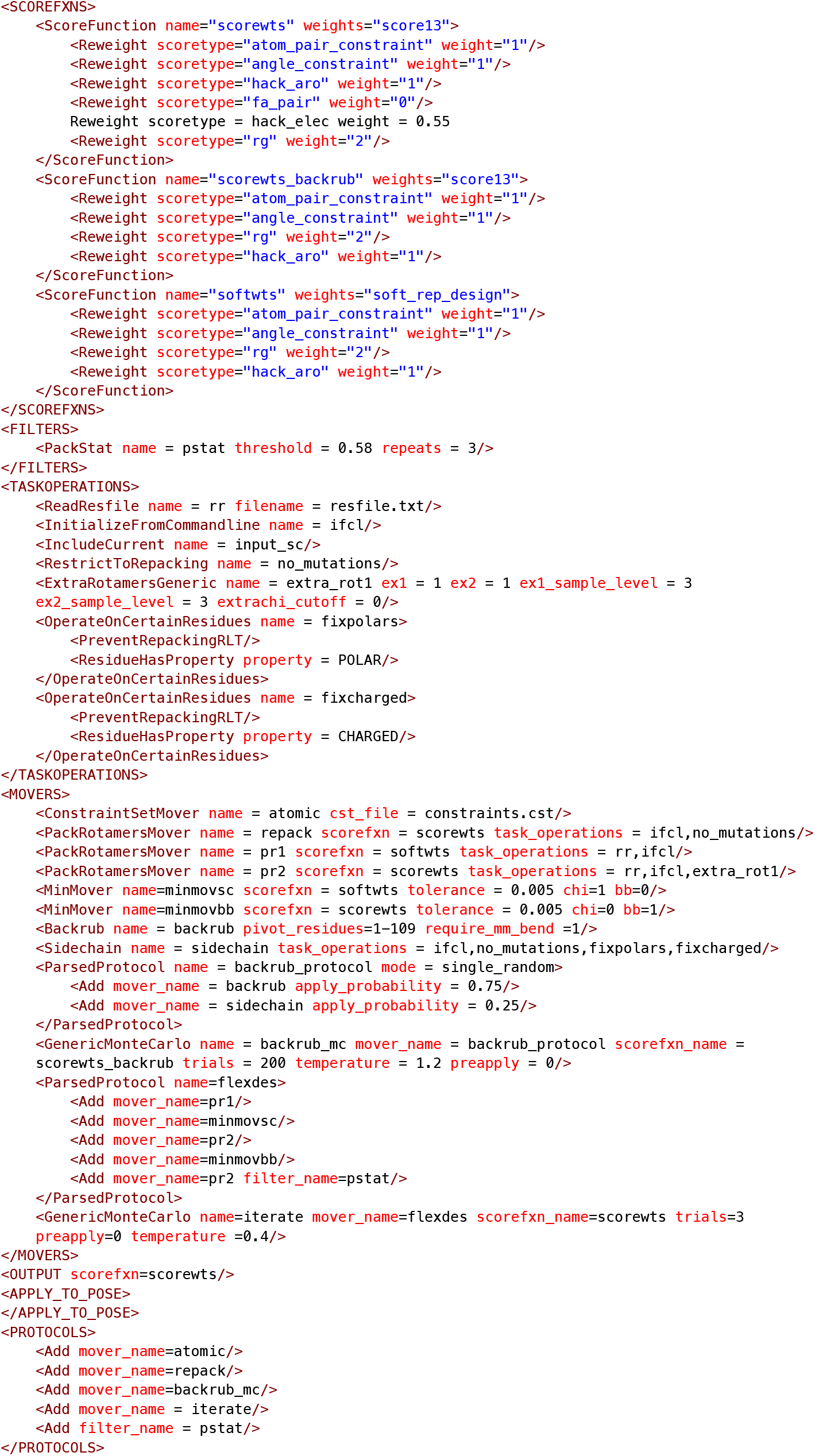

## Molecular Dynamics Input Files

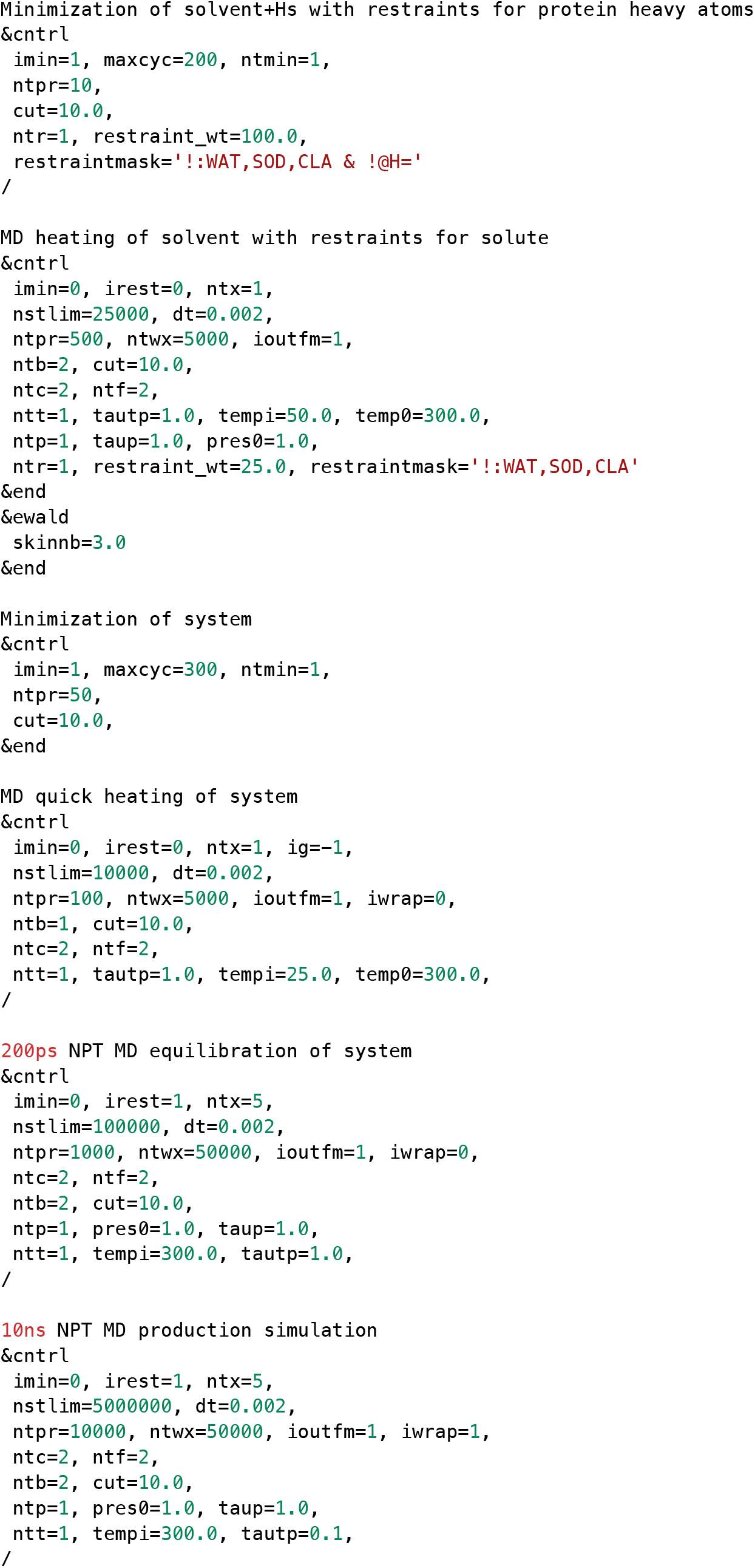

